# Concentration fluctuations due to size-dependent gene expression and cell-size control mechanisms

**DOI:** 10.1101/2021.10.18.464773

**Authors:** Chen Jia, Abhyudai Singh, Ramon Grima

## Abstract

Intracellular reaction rates depend on concentrations and hence their levels are often regulated. However classical models of stochastic gene expression lack a cell size description and cannot be used to predict noise in concentrations. Here, we construct a model of gene product dynamics that includes a description of cell growth, cell division, size-dependent gene expression, gene dosage compensation, and size control mechanisms that can vary with the cell cycle phase. We obtain expressions for the approximate distributions and power spectra of concentration fluctuations which lead to insight into the emergence of concentration homeostasis. Furthermore, we find that (i) the conditions necessary to suppress cell division-induced concentration oscillations are difficult to achieve; (ii) mRNA concentration and number distributions can have different number of modes; (iii) certain size control strategies are ideal because they maintain constant mean concentrations whilst minimising concentration noise. Predictions are confirmed using lineage data for *E. coli*, fission yeast and budding yeast.

## Introduction

Classical models of stochastic gene expression describe the fluctuations of protein and mRNA numbers in single cells and tissues [1–3]. These models have been successful at fitting experimental copy number distributions to estimate rate constants [4, 5]. They also have provided insight into the sources of intracellular fluctuations [6, 7], in their control via feedback mechanisms [8–10] and in their exploitation to generate oscillations and multi-stable states [11, 12].

However the majority of these models cannot be used to explain observations based on measurements of the content of individual cells collected over many generations [13] because the models ignore basic biological properties. For example in classical models, the mRNA and protein numbers will approach steady-state, while in reality, there is a regular pattern consisting of the increase of the molecule numbers during the cell cycle followed by their approximate halving at cell division. This pattern is strongly influenced by various sources of noise including cell cycle duration variability, DNA replication, and binomial partitioning at cell division. Some recent models [14–20] capture these phenomena and have led to a more detailed quantitative picture of gene expression; however they do not have an explicit cell size description. It has recently been shown [20] that a subset of these models which capture cell cycle duration variability via the Erlang distribution implicitly capture gene product number fluctuations induced by cell size fluctuations because the Erlang distribution can be expressed in terms of the parameters of any cell-size control mechanism [21]; however this is only the case when transcriptional activity of a promoter is independent of cell size, as e.g. found in *E. coli* [22].

In contrast, in yeast and mammalian cells, it has been shown that the initiation rate increases with cellular volume [23, 24]. This property has been hypothesized to be crucial to explaining concentration homeostasis, i.e. the observation that concentrations of many mRNAs and proteins vary little with cellular volume [25]. To capture this phenomenon there is the need of a model of gene expression that explicitly describes cell size; this can then be used to understand the factors giving rise to concentration homeostasis.

Here we develop a model that takes into account experimental observations of size-dependent gene expression, gene dosage compensation, DNA replication, cell division, and cell-size control mechanisms that vary according to cell cycle phases. By solving the model, we obtain insight into the statistical properties of concentration fluctuations in single cells across their cell cycle, the emergence of concentration homeostasis, and the non-trivial relationship between size control mechanisms and gene expression. Some of the predictions are confirmed by analysis of published data and others provide motivation for future experiments.

## Results

### Building a computational model that captures salient features of the underlying molecular biology

We consider a biologically detailed, stochastic model of gene expression coupled to cell size dynamics with the following properties (Fig. 1(a)).

**Figure 1:**
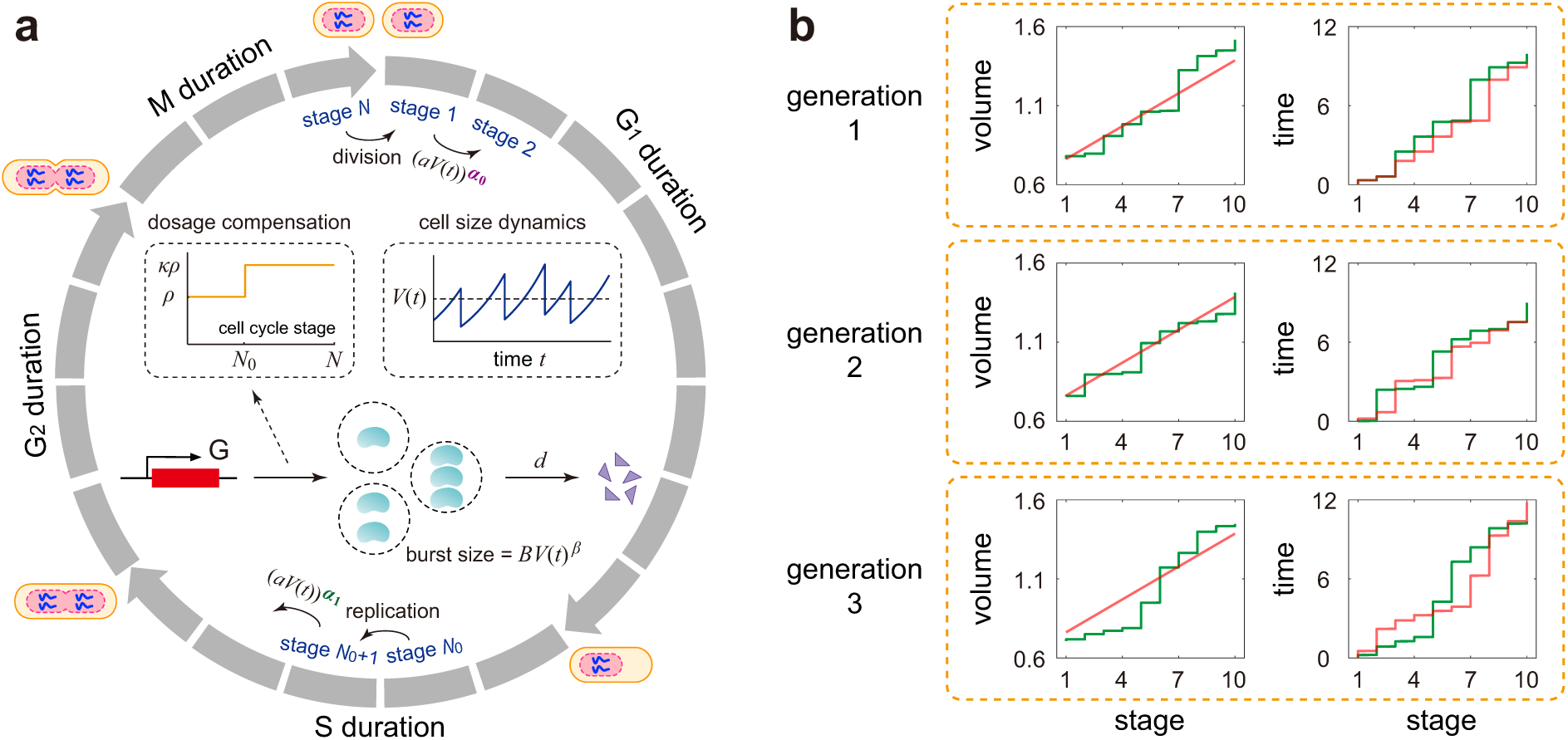
Model and its mean-field approximation. **(a)** A detailed stochastic model with both cell size and gene expression descriptions. The model consists of *N* effective cell cycle stages, gene replication at stage *N*_0_, cell division at stage *N*, bursty production of the gene product, and degradation of the gene product. The growth of cell volume is assumed to be exponential and gene dosage compensation is induced by a change in the burst frequency at replication from *ρ* to *κρ* with *κ* ∈ [1, 2) (see inset graphs). Both the transition rates between stages and the mean burst size depend on cell size in a power law form due to size homeostasis and balanced (or non-balanced) biosynthesis, respectively. The model also considers two-layer cell-size control with its strength varying from *α*_0_ to *α*_1_ upon replication. Recent studies have revealed that in many cell types, the accumulation of some activator to a critical threshold is used to promoter mitotic entry and trigger cell division, a strategy known as activator accumulation mechanism [26]. In *E. coli*, the activator was shown to be FtsZ [27–29]; in fission yeast, it was believed to be a protein upstream of Cdk1, the central mitotic regulator, such as Cdr2, Cdc25, and Cdc13 [30–33]. Biophysically, the *N* effective stages can be understood as different levels of the key activator. (**b**) Schematics of the changes in cell volume and time from birth as a function of cell cycle stage for the first three generations in the case of *N* = 10. In the full model (green curve), the gene expression dynamics is coupled with the following two types of fluctuations: noise in cell volume at each stage and noise in the time interval between two stages. The reduced model (red curve) ignores the former type of fluctuations by applying an mean-field approximation while retains the latter type of fluctuations. The reduced model approximates the full model excellently when *N* is large.

1. Cell volume grows exponentially with rate *g*. This assumption holds for a wide range of cell types [34–38].
2. Before replication, the synthesis of the gene product of interest, mRNA or protein, occurs at a rate *ρ* in bursts of a random size sampled from a geometric distribution with mean *B*′ [2]. In some types of bacteria, for some promoters, transcriptional activity is constant throughout the cell cycle which also implies the transcriptional parameters *ρ* and *B*′ are independent of cell size [22]. However, in yeast and mammalian cells, the products of many genes are produced in a balanced manner: the synthesis rate *ρB*′ is proportional to cell volume [23, 24]. Moreover, recent studies have shown that in the presence of balanced biosynthesis, cell size controls the synthesis rate via modulation of the burst size, instead of the burst frequency [23]. To unify results in various cell types, we assume that *B*′ = *BV* (*t*)^*β*^, where *V* (*t*) is cell volume, and *B* and 0 ≤ *β* ≤ 1 are two constants. Balanced (non-balanced) biosynthesis corresponds to the case of *β* = 1 (*β* = 0). We emphasize that while we assume bursty expression here, our model naturally covers constitutive expression since the latter can be regarded as a limit of the former when *ρ* → ∞, *B* → 0, and ⟨*n*⟩ ∝ *ρB* keeps constant [10, 39].
3. The gene product is degraded with rate *d* according to first-order kinetics.
4. The cell can exist in *N* effective cell cycle stages, denoted by 1, 2, …, *N*. Note that the *N* effective stages do not directly correspond to the four biological cell cycle phases (G_1_, S, G_2_, and M). Rather, each cell cycle phase is associated with multiple effective stages [40]. Recently, there has been evidence that yeast and mammalian cells may use different size control strategies before and after replication to achieve size homeostasis [41–44]. To model this effect, we assume that the transition rate from one stage to the next depends on cell volume in a power law form as 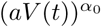 pre-replication and 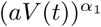 post-replication, where *a* is a constant and *α*_0_, *α*_1_ > 0 are the strengths of size control before and after replication. It is shown in Methods that *α*_0_ → 0, 1, ∞ correspond to mechanisms whereby the mean duration from birth to replication or the mean added size from birth to replication or the mean cell size at replication is independent of the size at birth, respectively. Similarly *α*_1_ → 0, 1, ∞ correspond to mechanisms whereby the mean duration from replication to division or the mean added size from replication to division or the mean cell size at division is independent of the size at replication, respectively. In other words, *α*_*i*_ → 0, 1, ∞ corresponds to timer, adder, and sizer, respectively. We define timer-like control to be when 0 < *α*_*i*_ < 1 and sizer-like control when 1 < *α*_*i*_ < ∞ [45].
5. Gene replication occurs when the cell transitions from a fixed stage *N*_0_ to the next. After replication, the number of gene copies doubles. If the burst frequency per gene copy does not change at replication, then the total burst frequency after replication is 2*ρ*. Dosage compensation is then modeled as a change in the burst frequency at replication from *ρ* to *κρ* with 1 ≤ *κ* < 2. [23, 46].
6. Cell division occurs when the cell transitions from stage *N* to stage 1. The volumes of the two daughter cells are assumed to be the same and exactly one half of the volume before division. Moreover, we assume that each gene product molecule has probability 1/2 of being allocated to each daughter cell.

### Simplifying the computational model via mean-field approximation

The model described above is very complex and hence it is no surprise that the master equation describing its stochastic dynamics is analytically intractable. The difficulty in solving the statistics of gene product fluctuations stems from the fact that they are coupled to two other types of fluctuations: (i) noise in the time elapsed between two cell cycle stages; (ii) noise in cell volume when a cell cycle stage is reached. Specifically, if at time *t* a cell enters stage *k* and has volume *V* (*t*), then it will jump to stage *k* + 1 after a random interval Δ*t* whose distribution is determined by the time-dependent rate function 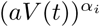. Since volume growth is exponential, the volume at stage *k* + 1 is given by *V* (*t* + Δ*t*) = *V* (*t*)*e*^*g*Δ*t*^. As can be seen from this reasoning, the fluctuations in cell volume at a certain stage depend on the fluctuations in the time to move to this stage from the previous stage.

To simplify this model, we devised a mean-field approximation on the volume dynamics, i.e. we ignore volume fluctuations at each stage but retain fluctuations in the time elapsed between two stages (Fig. 1(b)). Specifically, we calculate an approximate mean cell size at each stage by averaging over generations. This implies that the volume change from stage *k* to stage *k* + 1 is now deterministic and decoupled from the fluctuations in the time interval between the two stages. Within this approximation, the volume at division is twice that at birth implying that the mean cell cycle duration is *T* ≈ log(2)*/g* and thus the cell cycle frequency is *f* ≈ *g/* log(2). The time interval is itself exponentially distributed with rate 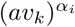, where *v*_*k*_ is the mean cell size at stage *k*. It is shown in Methods that the mean birth size *v*_1_ is approximately given by the solution of the implicit equation

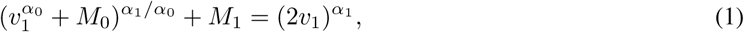

where 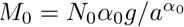 and 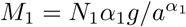. Moreover, the mean cell size at stage *k* is given by

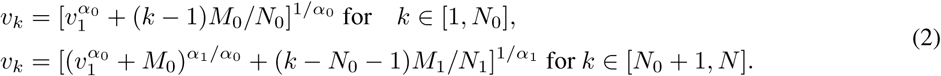

This approximation is generally valid provided the number of cell cycle stages *N* is large enough, which equivalently means that the variability in cell cycle duration is not very large [21]. In our model, the time from birth to replication occupies approximately a fixed proportion of the total cell cycle duration, with the proportion given by 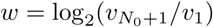. This is consistent with the stretched cell cycle model proposed in [47].

We now can construct a reduced model. Since we will be replacing a stochastic variable, the cell volume, by its mean, this is a type of mean-field approximation which has a long history of successful use in statistical physics [48]. In the reduced model, we choose the transition rate from stage *k* to the next to be 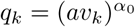 for *k* ∈ [1, *N*_0_] and 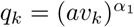 for *k* ∈ [*N*_0_ + 1, *N*], the mean burst size at stage *k* to be 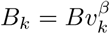, and the burst frequency at stage *k* to be *ρ*_*k*_ = *ρ* for *k* ∈ [1, *N*_0_] and *ρ*_*k*_ = *κρ* for *k* ∈ [*N*_0_ + 1, *N*] (Supplementary Fig. S1); these are the same rates as in the full model but with the instantaneous volume being replaced by its mean. The microstate of the reduced model can be represented by an ordered pair (*κ, n*), where *k* is the cell cycle stage and *n* is the copy number of the gene product. Let *p*_*k,n*_ denote the probability of observing microstate (*k, n*). The evolution of copy number dynamics is governed by a master equation which is given in Methods. A detailed comparison between stochastic simulations of the full and mean-field models show that they are in good agreement when *N* ≥ 15 and become practically indistinguishable when *N* ≥ 30 (Fig. 2 and Supplementary Fig. S2).

**Figure 2:**
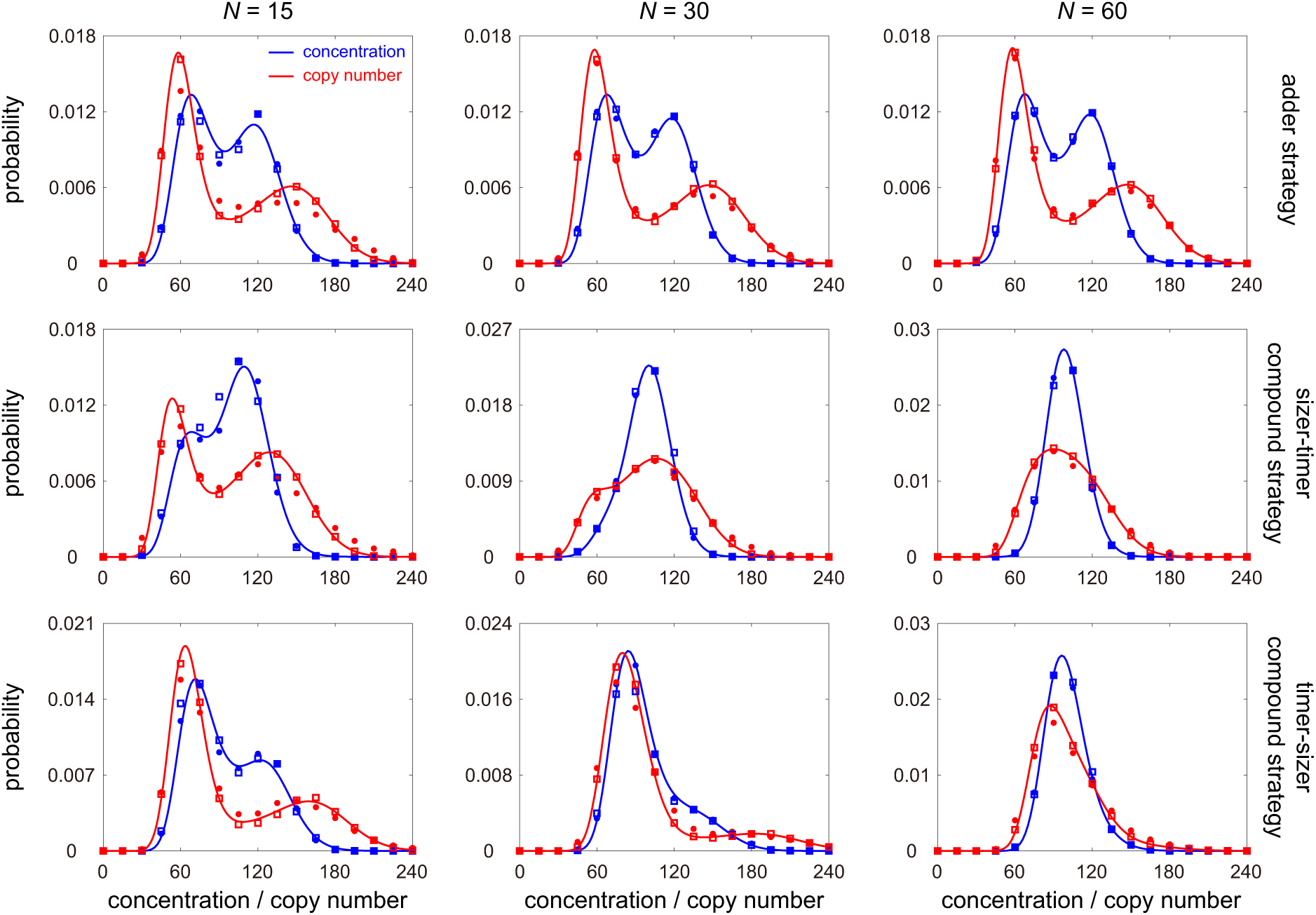
Comparison between the full and mean-field models under different choices of *N* and different size control strategies. The blue (red) dots show the simulated concentration (copy number) distribution for the full model obtained from the stochastic simulation algorithm (SSA). The blue (red) squares show the simulated concentration (copy number) distribution for the mean-field model obtained from SSA. The blue (red) curve shows the analytical approximate concentration (copy number) distribution given in Eq. (4) (Eq. (13)). The full and mean-field models are in good agreement when *N* ≥ 15 and they become almost indistinguishable when *N* ≥ 30. Moreover, the analytical solution performs well when *N* ≥ 15 and perform excellently when *N* ≥ 30. The model parameters are chosen as *N*_0_ = 0.4*N, B* = 1, *β* = 0, *κ* = 2, *d* = 1, *η* = 10. The growth rate *g* is determined so that *f* = 0.1. The parameters *ρ* and *a* are chosen so that the mean gene product number ⟨*n*⟩ = 100 and the mean cell volume ⟨*V* ⟩ = 1. The strengths of size control are chosen as *α*_0_ = *α*_1_ = 1 for the upper panel, *α*_0_ = 2, *α*_1_ = 0.5 for the middle panel, and *α*_0_ = 0.5, *α*_1_ = 2 for the lower panel. When performing SSA, the maximum simulation time for each lineage is chosen to be 10^5^. To deal with the time-dependent propensities in the full model, we use the numerical algorithm described in [49, Sec. 5].

### The concentration distribution is generally a sum of gamma distributions; for perfect concentration homeostasis, it is a single gamma

Many quantities of interest can be computed analytically using the master equation, Eq. (12), of the mean-field model. We then enforce cyclo-stationary conditions, i.e. steady state for the distribution of gene product number at each cell cycle stage. Let 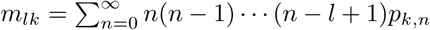 be the unnormalized *l*th factorial moment of copy numbers when the cell is at stage *k*. In particular, 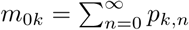 is the probability that the system is at stage *k*. For convenience, let *m*_*l*_ = (*m*_*lk*_) be the row vector composed of all *l*th factorial moments. At the steady state, *m*_0_ = (*m*_0*k*_) can be computed exactly as 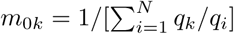 for single-cell lineage measurements, i.e. a single cell is tracked at a series of time points, and at division one of the two daughters is randomly chosen such that we have information of the stochastic dynamics along a cell lineage. Then the first two moments *m*_1_ and *m*_2_ are given by (Supplementary Note 1)

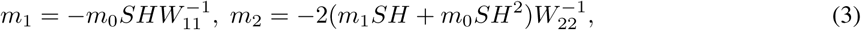

where *S* = diag(*ρ*_1_, *…, ρ*_*N*_), *H* = diag(*B*_1_, *…, B*_*N*_), and the matrices *W*_*ll*_, *l* ≥ 0 are given by

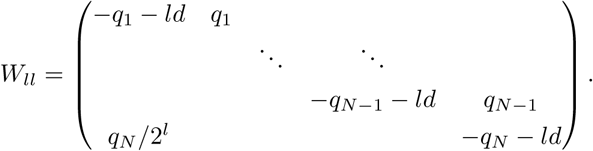

It follows that the concentration mean at stage *k* is given by *µ*_*k*_ = *m*_1*k*_/(*m*_0*k*_*v*_*k*_) and the variance at stage *k* is given by 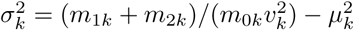. Once *µ*_*k*_ and 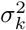 are known, the steady-state distribution of concentration can be computed approximately as (Methods)

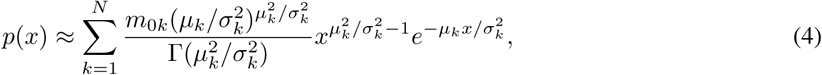

which is a mixture of *N* gamma distributions; the distribution of copy number can also be computed and is given in Methods. The analytical distribution agrees well with stochastic simulations when *N* ≥ 15 (Fig. 2 and Supplementary Fig. S2). The moments and distributions can also be computed for population measurements, i.e. all single cells in a population are measured at a particular time such that we have information of the stochastic dynamics across a growing population (Methods). For the rest of this paper, we focus on lineage calculations because the results from population calculations are qualitatively the same.

It can be shown that in the special case of balanced biosynthesis (*β* = 1, i.e. the mean burst size scales linearly with cell size) and perfect dosage compensation (*κ* = 1, i.e. the total burst frequency is independent of the number of gene copies), both the mean *µ*_*k*_ and variance 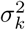 of concentration fluctuations are independent of stage *k*, i.e. invariant across the cell cycle, provided the number of gene product molecules is large (Supplementary Note 2). We shall call this condition perfect concentration homeostasis. In this case, we have *µ*_*k*_ = *ρB/d*_eff_ and 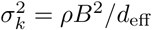, where *d*_eff_ = *d* + log(2)*f* is the effective decay rate of the gene product due to degradation and dilution at cell division. As well, in this case, Eq. (4) reduces to a gamma distribution and the theory agrees with the classical model of Friedman et al. [50] which has no explicit cell size description.

We note that the condition of perfect concentration homeostasis is stronger than what is normally referred to as concentration homeostasis which is indicated by an approximate linear relationship between the mean molecule numbers and cell volume [24, 25] — this ensures invariant mean concentrations but not a specific constraint on the variance of concentration fluctuations. The conditions necessary for perfect homeostasis do not generally exist in nature, e.g. the burst frequency per gene copy does not precisely halve upon replication and hence gene dosage compensation is never perfect [23, 46]. This means that generally some deviation of the concentration distribution from gamma is expected, and this is captured by Eq. (4).

### Accurate concentration homeostasis requires a fine tuning of dosage compensation and balanced biosynthesis

In what follows, we shall stick to the standard definition of concentration homeostasis in the literature [51], which does not specifically impose constraints on the variance but only requires constancy of the mean concentration across the cell cycle. In keeping with this definition, we introduce a new parameter *γ* as a measure of the accuracy of concentration homeostasis:

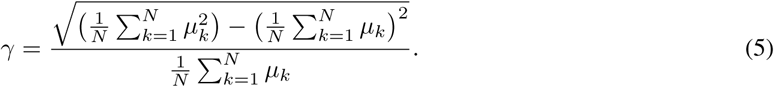

This is the coefficient of variation (CV) of the mean concentrations across all stages. The smaller the value of *γ*, the stronger the homeostatic mechanism.

We then use *γ* to investigate the conditions for accurate concentration homeostasis. In Fig. 3(a), we illustrate *γ* as a function of *β* and *κ*. Of particular interest is that there is a region of parameter space (shown in blue) where *γ* is minimised; this is approximately given by the curve 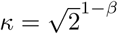 (shown by the yellow dash line). This implies that to maintain accurate homeostasis, a lack of balanced biosynthesis requires also a lower degree of dosage compensation.

**Figure 3:**
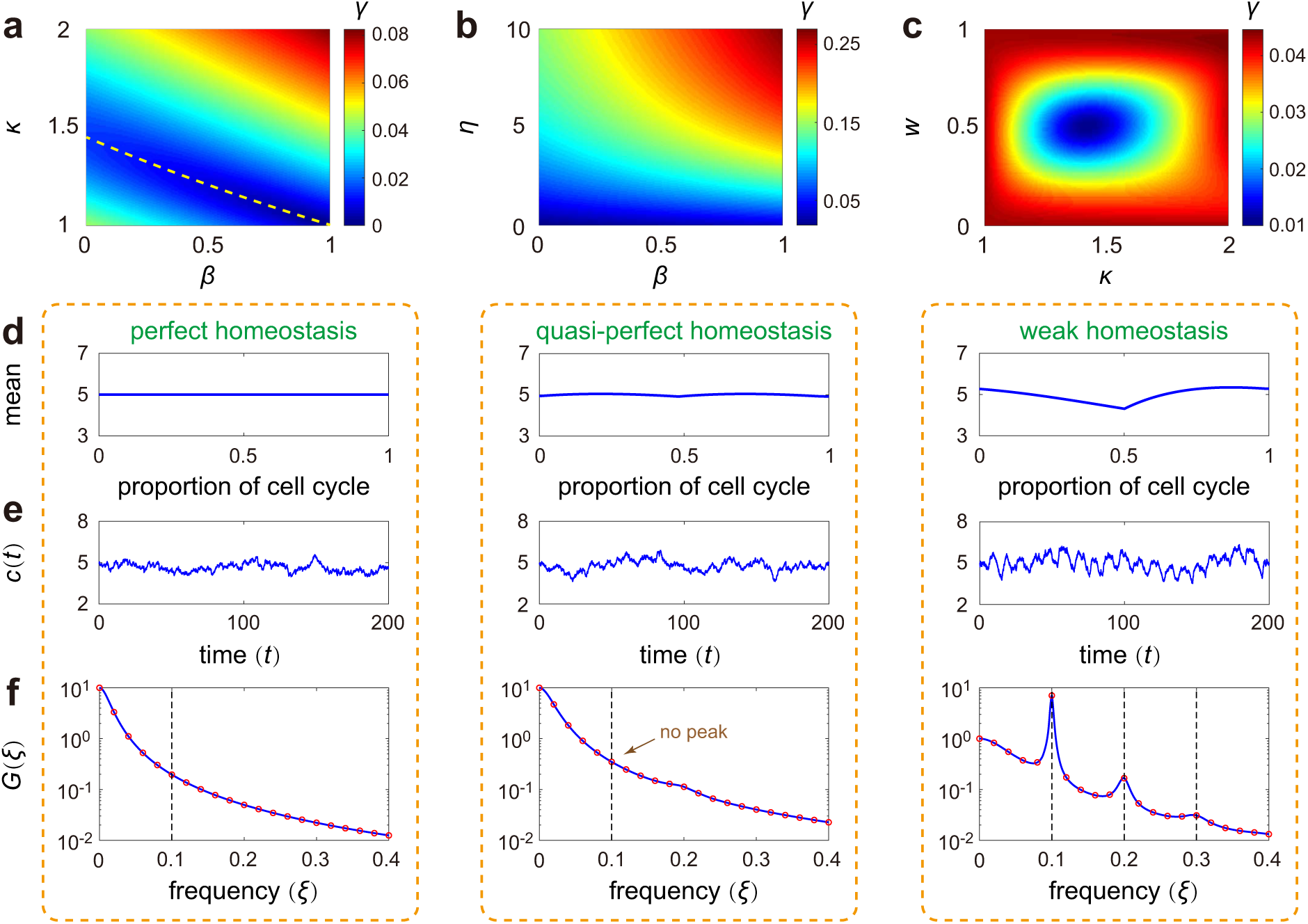
Concentration homeostasis and its relation to concentration oscillations. (**a**) Heat plot of *γ* (the accuracy of concentration homeostasis) versus *β* (the degree of balanced biosynthesis) and *κ* (the strength of dosage compensation). The yellow dashed line shows the exponential curve 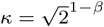, around which homeostasis is accurate. (**b**) Heat plot of *γ* versus *β* and *η* (the stability of the gene product) when there is no dosage compensation (*κ* = 2). Stable gene products give rise to more accurate homeostasis than unstable ones. (**c**) Heat plot of *γ* versus *κ* and *w* (the proportion of cell cycle before replication) when synthesis is non-balanced (*β* = 0). Homeostasis is the most accurate when 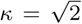 and *w* = 0.5. (**d**) The mean concentration *µ*_*k*_ in cell cycle stage *k* versus the proportion of cell cycle before stage *k*, which can be computed explicitly as *w*_*k*_ = log_2_(*v*_*k*_*/v*_1_). (**e**) Typical time traces of concentration. Here *c*(*t*) denotes the concentration at time *t* for a single cell lineage. (**f**) Theoretical (blue curve) and simulated (red circles) power spectra of concentration fluctuations. Throughout the paper, we normalize the spectrum so that *G*(0) = 1. Here the power spectrum 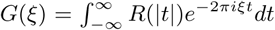 is defined as the Fourier transform of the autocorrelation function *R*(*t*). The theoretical spectrum is computed from Eq. (6), while the simulated spectrum is obtained by means of the Wiener-Khinchin theorem, which states that *G(ξ)* = lim_*T* → ∞_ ⟨| *ĉ*_*T*_(*ξ*)|^2^⟩/*T*, where 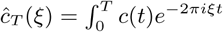 *dt* is the truncated Fourier transform of a single concentration trajectory over the interval [0, *T*] and the angled brackets denote the ensemble average over trajectories. The vertical lines show the cell cycle frequency *f* and its harmonics. See Supplementary Note 4 for the technical details of this figure.

Now we look in detail at the case where there is no dosage compensation (*κ* = 2). In Fig. 3(b), we illustrate *γ* as a function of *η* and *β*, where *η* = *d/f* is the ratio of the degradation rate to cell cycle frequency (a measure of the stability of the gene product). Note that *γ* becomes smaller as *η* decreases, implying that stable gene products (e.g. most proteins) give rise to better concentration homeostasis than unstable ones (e.g. most mRNAs). The combined results from Fig. 3(a),(b) indicate that homeostasis is the weakest when *β* = 1, *κ* = 2, and *η* is large. This shows that balanced biosynthesis, weak dosage compensation, and large degradation rates together can break homeostasis.

Next we look in detail at the case where synthesis is non-balanced (*β* = 0). In Fig. 3(c), we illustrate *γ* as a function of *κ* and *w*. Interestingly, we find that *γ* is remarkably small when 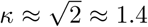 and *w* ≈ 0.5. The optimal values of *κ* and *w* are stable and depend very less on other parameters. This shows that when *β* = 0, accurate homeostasis can still be obtained at an intermediate strength of dosage compensation when replication occurs halfway through the cell cycle.

### Precise determination of the onset of concentration homeostasis using the power spectrum of fluctuations

Generally the time traces of gene product numbers increase from birth up till division time, at which point they halve. This periodicity would be also expected to some extent in concentrations. However clearly if there is strong concentration homeostasis, no such periodic behaviour would be visible. Hence it stands to reason that a lack of a peak in the power spectrum of concentration oscillations (computed from a single lineage) can be interpreted as strong concentration homeostasis. Stochastic simulations, shown in Fig. 3(d)-(f), indicate that this intuition is indeed the case: as *γ* increases (from left to right), the trajectory becomes more oscillatory and the power spectrum switches from monotonic to a peaked one at a non-zero value of frequency.

We next study the power spectrum analytically. Because the propensities of the reactions in our model are linear in gene product numbers, the autocorrelation of concentration fluctuations can be computed in closed form as (Supplementary Note 3)

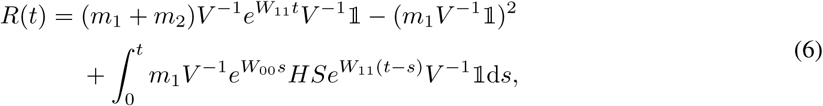

where *V* = diag(*v*_1_, *…, v*_*N*_) and 𝟙 = (1, *…*, 1)^*T*^. The power spectrum is then the Fourier transform of the autocorrelation which can be computed straightforwardly. It can be shown that the power spectrum is a weighted sum of *N* Lorentzian functions, which either decreases monotonically or has an off-zero peak around the cell cycle frequency *f*, whose height is proportional to both *N* ^2^ and (Supplementary Note 3)

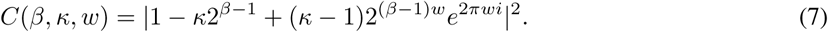

Furthermore, when *N* ≫ 1, the power spectrum also has peaks around the higher order harmonics of *f* (right panel of Fig. 3(f)).

Generally, the condition *C*(*β, κ, w*) = 0 is the necessary condition for the power spectrum to lack an off-zero peak which means that strong homeostasis is present. This can be achieved when *β* = *κ* = 1 (balanced biosynthesis and perfect dosage compensation) which implies also *γ* = 0. However interestingly, it can be achieved also when balanced biosynthesis is broken. When *β* ≠ 1, Eq. (7) vanishes when 1 − *κ*2^*β*−1^ + (*κ* − 1)2^(*β*−1)*w*^ cos(2*πw*) = 0 and sin(2*πw*) = 0, i.e. 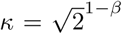 and *w* = 0.5. In this case, *γ* ≠0 and thus according to Eq. (5), it means that there is still some variation of the mean concentration across the cell cycle. Hence we refer to this case as quasi-perfect homeostasis (middle column of Fig. 3(d)-(f)). This also corresponds to the minimum shown by the yellow dashed line in Fig. 3(a). In particular, when *β* = 0, quasi-perfect homeostasis is obtained when 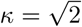 and *w* = 0.5 (Fig. 3(c)). The optimal value of 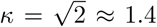 is comparable to the experimental values of *κ* measured for *Oct4* and *Nanog* genes in mouse embryonic stem cells [46]. However generally the conditions for strong homeostasis, i.e. no peak in the power spectrum, appear quite restrictive (*β* = *κ* = 1 or *β* ≠1, 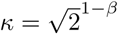, and *w* = 0.5) and hence likely this is rarely seen in nature.

It has been shown [20] that for unstable gene products and non-balanced biosynthesis, the time traces of copy number display the strongest oscillation regularity when *w* = 0.5; here we have shown that the time traces of concentration display the weakest regularity when *w* = 0.5 (Supplementary Fig. S3). This shows that copy number and concentration time traces may exhibit totally different dynamical behaviors.

### A sizer-timer size control strategy maintains concentration homeostasis while minimising concentration noise

Recently, evidence has emerged that budding yeast, fission yeast, and mammalian cells may exhibit multiple layers of size control in each cell cycle, e.g. there can be different size control strategies at work before and after replication [41–44]. A natural question is why eukaryotic cells apply such two-layer control strategies to achieve size homeostasis. In our model, we assume that the size control strength varies from *α*_0_ to *α*_1_ upon replication. It can be shown that when the variability in cell size is small, the birth size *V*_*b*_ and the division size *V*_*d*_ are interconnected by *V*_*d*_ = 2^1−*α*^*V*_*b*_ + *ϵ*, where *α* = *wα*_0_ + (1 − *w*)*α*_1_ is the effective control strength over the whole cell cycle and *ϵ* is a noise term independent of *V*_*b*_ (Methods). The cell behaves as an overall timer/adder/sizer when *α* → 0, 1, ∞. Note that in real systems, *α* is never much larger than 1, e.g. for fission yeast, which is known to have a sizer-like mechanism, it has recently been estimated that *α* is 1.4 − 2.1 [52].

To understand the effect of size control on concentration homeostasis, we illustrate *γ* as a function of *β, α*_0_, and *α*_1_ (Fig. 4(a),(b)). Recall that balanced biosynthesis, weak dosage compensation, and unstable gene products may break concentration homeostasis. When synthesis is balanced (*β* = 1) and dosage compensation is not very strong (*κ* is not very close to 1), we find that homeostasis depends strongly on size control and its accuracy is significantly enhanced if cells use different size control strategies before and after replication (see also Supplementary Fig. S4(a),(b)). Homeostasis is strong if (i) the cell behaves as a sizer before replication and a timer after replication or (ii) the cell behaves as a timer before replication and a sizer after replication. Thus such sizer-timer or timer-sizer compound control strategies may help eukaryotic cells maintain accurate homeostasis without any clearly distinguishable concentration oscillations (Fig. 4(e)) for genes with a wide range of dynamic parameters. In contrast, when synthesis is non-balanced (*β* = 0) such as in bacteria, the same strategy before and after replication is ideal to maintain concentration homeostasis (Supplementary Fig. S4(e)-(f)).

**Figure 4:**
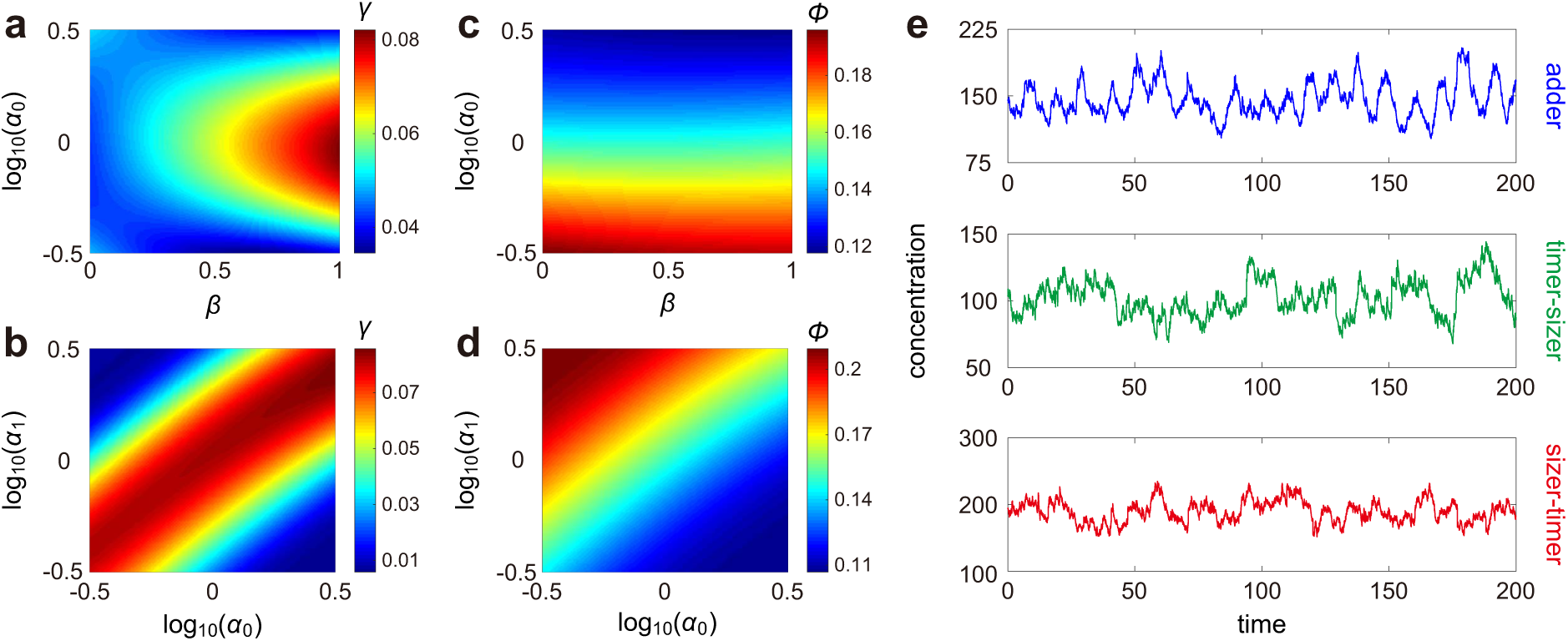
Influence of size control on concentration homeostasis and concentration noise. (**a**),(**b**) Heat plot of *γ* (the accuracy of concentration homeostasis) versus *β* (the degree of balanced biosynthesis), *α*_0_ (the strength of size control before replication), and *α*_1_ (the strength of size control after replication). When synthesis is balanced (*β* = 1), concentration homeostasis is accurate when the cell applies different size control strategies before and after replication. Both the timer-sizer and sizer-timer compound strategies enhance the accuracy of homeostasis. (**c**),(**d**) Heat plot of *ϕ* (noise in concentration) versus *β, α*_0_, and *α*_1_. While the timer-sizer compound strategy leads to accurate concentration homeostasis, it results in large gene expression noise. The sizer-timer compound strategy both maintains homeostasis and reduces noise. The model parameters are chosen as *N* = 50, *N*_0_ = 23, *ρ* = 1.7, *B* = 1, *κ* = 2, *d* = 0.1, *η* = 1 in (a)-(d), *α*_1_ = 1 in (a),(c), and *β* = 1 in (b),(d). The growth rate *g* is determined so that *f* = 0.1. The parameter *a* is chosen so that the mean cell volume ⟨*V*⟩ = 1. (**e**) Typical time traces of concentration under three different size control strategies: adder (upper panel), timer-sizer (middle panel), and sizer-timer (lower panel). See Supplementary Note 5 for the technical details of this figure.

Furthermore, we find that size control also affects gene expression noise. In our model, the steady-state concentration mean *µ* and variance *σ*^2^ can be computed analytically as 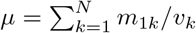 and 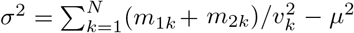. Noise in concentration can be characterized by *ϕ* = *σ*^2^*/µ*^2^, the CV squared of concentration fluctuations. Fig. 4(c),(d) illustrate *ϕ* as a function of *β, α*_0_, and *α*_1_, from which we can see that concentration noise depends only weakly on *β* but depends strongly on size control. Noise is minimised when the cell behaves as a sizer before replication and a timer after replication, and is maximized when the control strategy varies from timer to sizer at replication, whether synthesis is balanced or not (Fig. 4(d) and Supplementary Fig. S4). This suggests that while the timer-sizer compound strategy is conducive to concentration homeostasis, it gives rise to large concentration noise (Fig. 4(e)). However, the sizer-timer compound strategy both maintains homeostasis and reduces noise.

In a recent work [44], it was found that both the duration of the G_1_ phase and the added volume in G_1_ decrease with birth volume in two types of cancerous epithelial cell lines (HT29-hgem and HeLa-hgem), indicating that the control strategy before replication is sizer-like. Moreover, they also found that the duration of the S-G_2_ phase decreases and the added volume in S-G_2_ increases with the volume at the G_1_/S transition in both cell types, showing that the cell uses a timer-like strategy after replication. Similar results have been found in yeast [41–43]. Our results may explain why the sizer-timer compound strategy is preferred over others and hence commonly used in nature.

### Bimodality in the concentration distribution is rarer than bimodality in the distribution of molecule numbers

To gain a deeper insight into gene product fluctuations, we contrast the copy number and concentration distributions in Fig. 5(a)-(c). Clearly, the shapes of the two can be significantly different; for some parameter sets, one is unimodal and the other is bimodal. Both distributions can display bimodality, due to a change in the burst frequency upon replication; however, the concentration distribution is generally less wide than the copy number distribution, which agrees with experimental observations [25]. According to simulations, in the presence of balanced biosynthesis, the intrinsic noise in concentration is almost the same as that in copy number, while the extrinsic noise in concentration is always smaller than that in copy number (Supplementary Note 1 and Supplementary Fig. S5).

**Figure 5:**
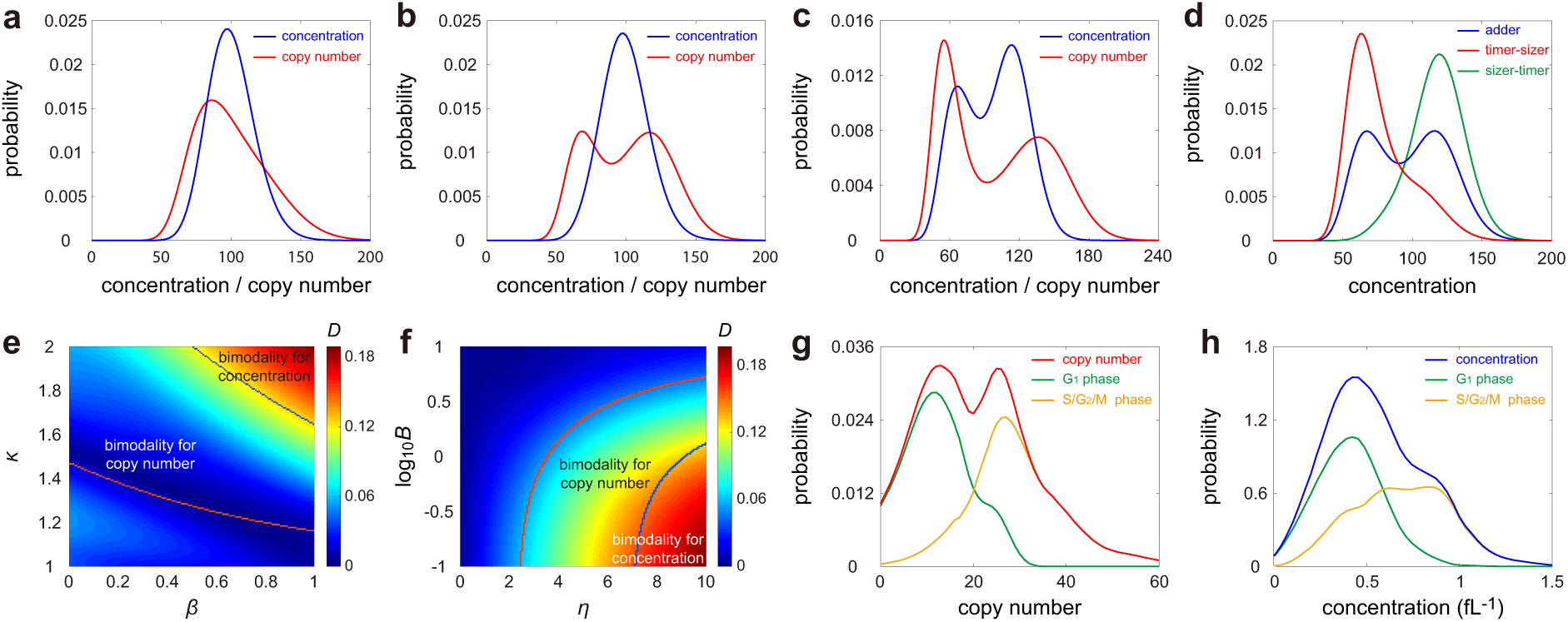
Influence of model parameters on copy number and concentration distributions. **(a)-(c)** The copy number (red curve) and concentration (blue curve) distributions display different shapes under different choices of parameters. (a) Both distributions are unimodal. (b) The concentration distribution is unimodal while the copy number one is bimodal. (c) Both distributions are bimodal. **(d)** Concentration distributions under three different size control strategies: adder (blue curve), timer-sizer (red curve), and sizer-timer (green curve). Two-layer control strategies lead to unimodal concentration distributions that are not far from gamma. (**e**),(**f**) Heat plot of the Hellinger distance *D* of the concentration distribution from its gamma approximation versus *β* (the degree of balanced biosynthesis), *κ* (the strength of dosage compensation), *η* (the stability of the gene product), and *B* (the mean burst size). The violet (blue) curve encloses the region for the copy number (concentration) distribution being bimodal. The bimodal region for the copy number distribution is much larger than that for the concentration distribution. (**g**),(**h**) The mRNA copy number (red curve) and concentration (blue curve) distributions for the *HTB2* gene in budding yeast measured using smFISH. The green (yellow) curves show the distributions of cells in the G_1_ (G_2_-S-M) phase. No apparent dosage compensation was observed with the mean copy number (concentration) in G_2_-S-M being 2.22 (1.66) times greater than that in G_1_. The data shown are published in [53]. See Supplementary Note 6 for the technical details of this figure.

Recall that concentration has the gamma distribution when *β* = *κ* = 1 and ⟨*n*⟩ ≫ 1. To understand the deviation from gamma, we depict the Hellinger distance *D* of the concentration distribution from its gamma approximation as a function of *β, κ, η*, and *B* (Fig. 5(e),(f)). Note that Fig. 5(e) is similar to Fig. 3(a), implying that the deviation from gamma is closely related to concentration homeostasis. Accurate homeostasis usually results in a concentration distribution that is gamma-shaped. In particular, stable gene products usually have a unimodal concentration distribution that is not far from gamma. Since sizer-timer and timer-sizer compound strategies enhance the accuracy of homeostasis, they both lead to a smaller deviation from gamma (Fig. 5(d)). In Fig. 5(e),(f) we show the regions of parameter space where the concentration and copy number distributions are unimodal and bimodal. Both distributions display bimodality when *β, κ*, and *η* are large and *B* is small. However, the region of parameter space for observing bimodality in copy number distributions is much larger than that for concentration distributions. Bimodality in the concentration distribution occurs only in the presence of balanced biosynthesis, weak dosage compensation, large degradation rates, and small burst size.

### Confirmation of theoretical predictions using bacterial and yeast data

Next we use published experimental data to test some of the predictions of our model: (i) unstable gene products can have a concentration distribution with either the same or a smaller number of modes than the copy number distribution; (ii) strong concentration homeostasis is rare, i.e. while the gene product numbers may scale approximately linearly with cell volume, nevertheless oscillations in concentration due to the periodicity of cell division are still detectable; (iii) deviations from perfect homeostasis result in deviations of the concentration distribution from gamma. We used three data sets reporting mRNA or protein measurements for bacteria, budding yeast, and fission yeast, to test these results, as follows.

To test prediction (i), we used the population data collected in diploid budding yeast cells using single-molecule mRNA fluorescence in situ hybridization (smFISH) [53]. In this data set, the mRNA abundance and cell volume are measured simultaneously for four genes: *HTB1, HTB2, HHF1*, and *ACT1*, with the cell cycle phase (G_1_, S, and G_2_/M) of each cell being labelled. These mRNAs are produced in a balanced manner [53] and expected to be unstable since the median mRNA half-life in budding yeast is ∼ 20 min [54], which is much less than the mean cell cycle duration of 1.5 − 2.5 hr [55, 56]. Interestingly, for *HTB2*, we observed an apparent bimodal distribution for mRNA number and a unimodal distribution for mRNA concentration (shown by the red and blue lines in Fig. 5(g),(h)). Calculating the distributions conditional on the G_1_ and S-G_2_-M phases (shown by the green and yellow lines) clarifies that the bimodality stems from replication and a lack of dosage compensation. Similar distribution shapes were also observed for *HTB1* and *ACT1*, although bimodality in the concentration distribution is significantly less apparent; for *HHF1*, both distributions are unimodal (Supplementary Fig. S6).

To test predictions (ii) and (iii), we used the lineage data collected in *E. coli* and haploid fission yeast cells using a mother machine [13, 57]. In the *E. coli* data set [13], the time course data of cell size and fluorescence intensity of a stable fluorescent protein were recorded for 279 cells lineages under three growth conditions. In the fission yeast data set [57], similar data for a stable fluorescent protein were monitored for 10500 cell lineages under seven growth conditions. For both cell types, we illustrate the change in the mean concentration across the cell cycle, as well as the distribution and power spectrum of concentration fluctuations, in Fig. 6(a)-(c) and (e)-(g). In agreement with prediction (ii), from Fig. 6(a),(b) we see that for *E. coli*, while the mean concentration is practically invariant across the cell cycle, nevertheless homeostasis is not strong enough to suppress concentration oscillations whose signature is a peak at a non-zero frequency in the power spectrum. A similar phenomenon was also observed for fission yeast (Fig. 6(e),(f)). The higher-order harmonics of concentration fluctuations (second and third peaks in the spectrum) can be seen for fission yeast but not for *E. coli*; this is likely due to a stronger periodicity (smaller variability) in cell cycle duration in fission yeast compared to that in *E. coli* [21, 52]. Finally, in agreement with prediction (iii), from Fig. 6(d),(h) we see that for both organisms, the Hellinger distance *D*, which measures the distance of the concentration distribution from a gamma, is positively correlated with the homeostasis accuracy *γ* and the off-zero peak height *H*, both of which measure how strong homeostasis is.

**Figure 6:**
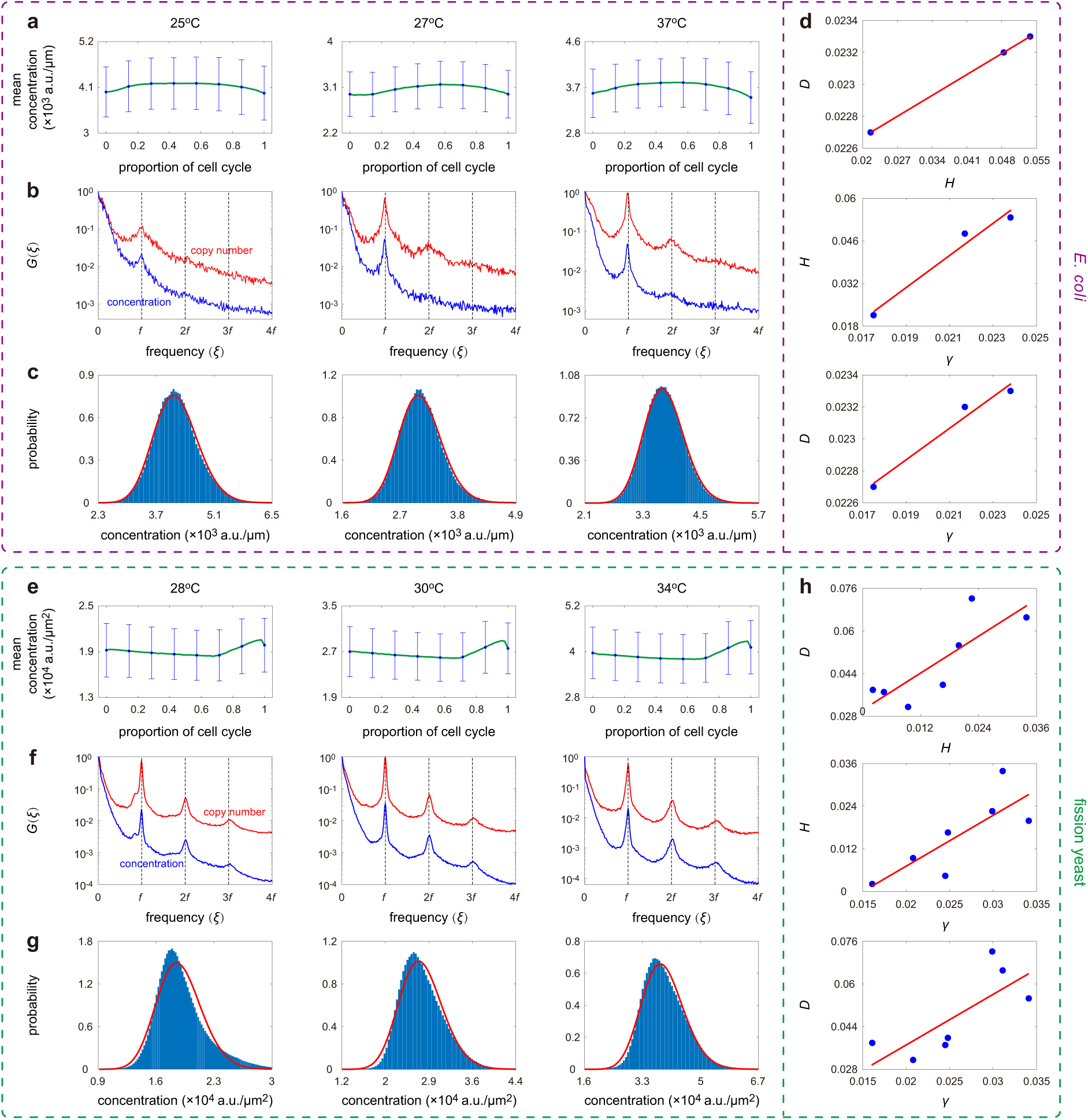
Analysis of single-cell lineage data of concentration fluctuations in *E. coli* and fission yeast. The data shown are published in [13, 57]. The *E. coli* data set contains the lineage measurements of both cell length and fluorescence intensity of a stable fluorescent protein at three different temperatures (25°C, 27°C, and 37°C). The fission yeast data set contains the lineage measurements of both cell area and fluorescence intensity of a stable fluorescent protein under seven different growth conditions (Edinburgh minimal medium (EMM) at 28°C, 30°C, 32°C, and 34°C and yeast extract medium (YE) at 28°C, 30°C, and 34°C). (**a**) The change in the mean concentration across the cell cycle for *E. coli* under three growth conditions. Here, we divide each cell cycle into 50 pieces and compute the mean and standard deviation (see error bars) of all data points that fall into each piece. The CV of the mean concentrations of all pieces gives an estimation of the homeostasis accuracy *γ*. (**b**) Power spectra of copy number (red curve) and concentration (blue curve) fluctuations under three growth conditions. (**c**) Experimental concentration distributions (blue bars) and their gamma approximations (red curve) under three growth conditions. (**d**) Scatter plots between *γ* (the accuracy of concentration homeostasis), *H* (the height of the off-zero peak of the power spectrum), and *D* (the Hellinger distance between the concentration distribution from its gamma approximation) under three growth conditions. (**e**)-(**h**) Same as (a)-(d) but for fission yeast. Here (e)-(g) only show the three growth conditions for YE, while (h) shows the scatter plots between *γ, H*, and *D* under all seven growth conditions (for both EMM and YE). Both cell types show remarkable positive correlations between *γ, H*, and *D*.

## Discussion

In this paper, we have developed and solved a complex model of gene product fluctuations which presents a unified description of intracellular processes controlling cell size, cell cycle phase, and their coupling to gene expression. The analytical difficulties in solving a model with considerable biological details were circumvented by means of a novel type of mean-field approximation which involves neglecting cell size fluctuations such that the cell size at a given fraction of the cell cycle is independent of the generation number. This approximation led to a reduced model from which we derived expressions for the moments, distributions, and power spectra of both gene product number and concentration fluctuations. We showed by simulations that the full and reduced models are in good agreement over all of parameter space provided the number of cell cycle stages *N* is greater than 15. Previous work estimated *N* to be 15 − 38 for the bacterium *E. coli*, according to temperature [21], 16 − 50 for fission yeast, depending on temperature and medium [52], 27 for human B-cells, 21 for rat 1 fibroblasts, 50 for human mammary epithelial cells, and 59 for the cyanobacterium *S. elongatus* [18].

We note that whilst the majority of gene expression models have no cell size description, recently a few pioneering papers have considered extensions to include such a description. In [51], it was shown that concentration homeostasis can arise when RNA polymerases are limiting for transcription and ribosomes are limiting for translation. This model takes into account gene replication and cell division but neglects dosage compensation and multi-layer size control, and some of the mechanistic details are specific to *E. coli*. In addition, the analysis is deterministic and while stochastic simulations are presented, a detailed study of fluctuations in molecule number and concentration is missing. In [58], mRNA number distributions are approximated by a negative binomial using a model that takes into account replication, division, dosage compensation, and balanced biosynthesis. The main limitations of this approach is its lack of flexibility to capture complex distributions and the assumption that the size control strategy is timer. In [59], a simplified model is considered whereby replication, dosage compensation, and multi-layer size control are neglected (an adder is the focus of the paper). A stochastic analysis is presented whereby the linear noise approximation (LNA) [60, 61] is extended to obtain the first two moments of copy number and concentration fluctuations. The focus of the study was to shed light on the similarities and differences with classical models that are in widespread use. Because the use of the LNA, only a Gaussian approximation to the copy number and concentration distributions are possible. Our model differs from the above models in numerous ways, by (i) including a wider description of relevant biological processes; (ii) capturing complex distributions and power spectra of concentration fluctuations using a novel non-LNA-based approach; (iii) showing that accurate homeostasis results from a complex tradeoff between dosage compensation, balanced biosynthesis, degradation, and the timing of replication; (iv) showing that concentration homeostasis is typically not strong enough to suppress concentration oscillations due to cell division; (v) showing that the number of modes of mRNA number distributions is greater than or equal to that of mRNA concentration distributions; (vi) showing that a sizer-timer compound strategy is ideal because it maintains accurate concentration homeostasis whilst minimising concentration noise.

Among the limitations of our model are the assumption that the parameters *β* and *κ* are independent of each other (in reality the two are likely related because both balanced biosynthesis and dosage compensation emerge from RNA polymerase dynamics), and its lack of a description of growth rate variability [44, 62, 63] and of regulation via feedback mechanisms [10, 12, 64–68]. In particular regarding the latter, it remains to be seen how the plethora of results on noise-induced bifurcations and oscillations derived using classical models are modified when size-dependent transcription and cell-size control strategies are introduced.

## Methods

### Derivation of the mean cell volume at each stage using a mean-field approximation

Let *V*_*b*_, *V*_*r*_, and *V*_*d*_ denote the cell volumes at birth, replication, and division, respectively. Let *α*_0_ and *α*_1_ denote the strengths of size control before and after replication, respectively. Here we shall prove that (i) the generalized added size before replication,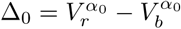, has an Erlang distribution with shape parameter *N*_0_ and mean 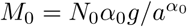 and (ii) the generalized added size after replication, 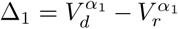, is also Erlang distributed with shape parameter *N*_1_ = *N* − *N*_0_ and mean 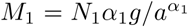. Using this property, *N*_0_ and *N*_1_ can be experimentally inferred as the inverse CV squared of Δ_0_ and Δ_1_, respectively.

To prove this, we first focus on the volume dynamics before replication. Note that when *V*_*b*_ is fixed, the cell volume at time *t* is given by 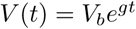. Since the transition rate from one stage to the next is equal to 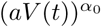 before replication, the distribution of the transition time *T* is given by

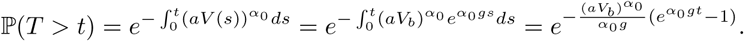

This shows that

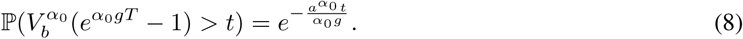

Let *v*_*k*_ be the volume when the cell first enters stage *k* with *v*_1_ = *V*_*b*_ and 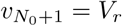. Note that 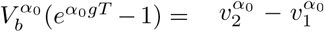 is the generalized added size at stage 1. Hence Eq. (8) suggests that 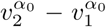 is exponentially distributed with mean 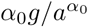. Similarly, we can prove that the generalized added size 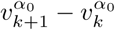 at stage *k* is also exponentially distributed with mean 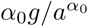 for each *k* ∈ [1, *N*_0_]. As a result, the generalized added size before replication, 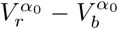, is the independent sum of *N*_0_ exponentially distributed random variables with the same mean 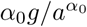. This indicates that 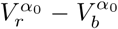 has an Erlang distribution with shape parameter *N*_0_ and mean 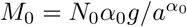, and is also independent of *V*_*b*_. The fact that 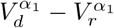 is Erlang distributed with shape parameter *N*_1_ and mean *M*_1_, and is also independent of both *V*_*b*_ and *V*_*r*_ can be proved in the same way.

Moreover, it is easy to see that for each *k* ∈ [1, *N*_0_], the generalized added size 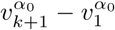, has an Erlang distribution with shape parameter *k* and mean 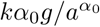. Similarly, for each *k* ∈ [*N*_0_ + 1, *N*], the generalized added size 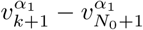 is also Erlang distributed with shape parameter *k* − *N*_0_ and mean 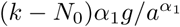.

We next focus on three important special cases. When *α*_0_ → 0, the transition rate from stage *k* ∈ [1, *N*_0_] to the next is a constant independent of *k* and thus the time elapsed from birth to replication has an Erlang distribution independent of the birth size; this corresponds to the timer strategy. When *α*_0_ = 1, the added size *V*_*r*_ − *V*_*b*_ before replication has an Erlang distribution independent of the birth size; this corresponds to the adder strategy. When *α*_0_ → ∞, the *α*_0_th power of the cell size at replication, 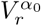, has an Erlang distribution independent of the birth size; this corresponds to the sizer strategy. Similarly, when *α*_1_ → 0, the transition rate from stage *k* ∈ [*N*_0_ + 1, *N*] to the next is a constant independent of *k* and thus the time elapsed from replication to division has an Erlang distribution independent of the replication size; this corresponds to the timer strategy. When *α*_1_ = 1, the added size *V*_*d*_ − *V*_*r*_ after replication has an Erlang distribution independent of the replication size; this corresponds to the adder strategy. When *α*_1_ → ∞, the *α*_1_th power of the cell size at division, 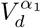, has an Erlang distribution independent of the replication size; this corresponds to the sizer strategy.

Since the exact mean cell volume at each stage is difficult to obtain, we make the approximation that the number of stages is large, which holds for most cell types and most growth conditions [18, 20, 21]. When *N* ≫ 1, the fluctuations in cell volume at each stage are very small and thus can be ignored. In this case, both Δ_0_ = *M*_0_ and Δ_1_ = *M*_1_ can be viewed as deterministic and *V*_*d*_ = 2*V*_*b*_. Thus the typical birth size *V*_*b*_ is the solution of the implicit equation

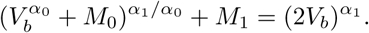

Moreover, when *N* is large, both 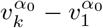 and 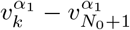 can be viewed as deterministic and replaced by their means, as given above. Thus the typical cell size at stage *k* is given by

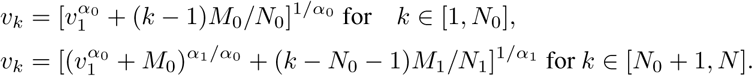

### Effective size control strategy over the whole cell cycle

In our model, we assume that the size control strategies before and after replication can be different with the strength of size control changing from *α*_0_ to *α*_1_ at replication. The following question naturally arises: what is the size control strategy over the whole cell cycle? To answer this, we first focus on the cell size dynamics before replication. Recall that the generalized added size 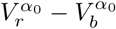 before replication has an Erlang distribution and is independent of *V*_*b*_. When the fluctuations of *V*_*b*_ and *V*_*r*_ are small, we have the following Taylor expansions:

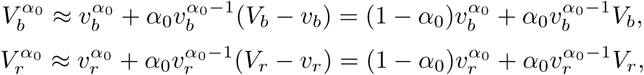

where *v*_*b*_ = *v*_1_ and 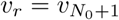 are the typical values of *V*_*b*_ and *V*_*r*_, respectively. Therefore, we obtain

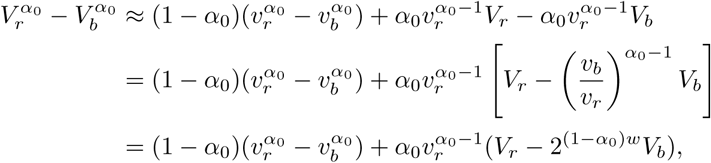

where *w* = log_2_(*v*_*r*_*/v*_*b*_) is the proportion of cell cycle before replication. This shows that

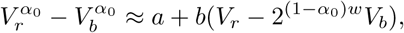

where 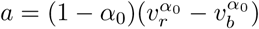 and 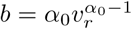 are two constants. Hence 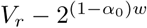*V*_*b*_ is approximately independent of *V*_*b*_, i.e.

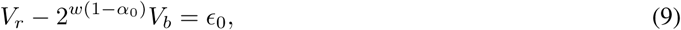

where *ϵ*_0_ is a noise term independent of *V*_*b*_ and which has a mean greater than zero. Similarly, for the cell size dynamics after replication, we have

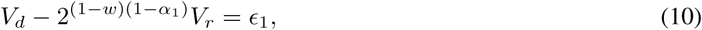

where *ϵ*_1_ is a noise term independent of both *V*_*b*_ and *V*_*r*_, and which has a mean greater than zero. Combining the above two equations, we can eliminate *V*_*r*_ and obtain

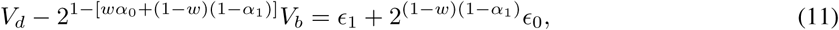

where the right-hand side is a noise term independent of *V*_*b*_. Eqs (9) and (10) again show that *α*_*i*_ → 0, 1, ∞ corresponds to timer, adder, and sizer, respectively.

To proceed, note that if the strength of size control does not change upon replication, i.e. *α*_0_ = *α*_1_ = *α*, then similar reasoning suggests that *V*_*d*_ − 2^1−*α*^*V*_*b*_ is a noise term independent of *V*_*b*_. Therefore, if the strength of size control changes from *α*_0_ to *α*_1_ at replication, then it follows from Eq. (11) that the effective size control strength *α* over the whole cell cycle should be defined as

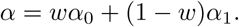

The cell behaves as an overall timer/adder/sizer when *α* → 0, 1, ∞.

The above discussion also provides a method of inferring *α*_0_ and *α*_1_ from experimental data. From Eq. (9), the slope of the regression line through the data points (*V*_*b*_, *V*_*r*_) gives an estimate of (1 − *α*_0_)*w*, from which *α*_0_ can be inferred. Similarly, from Eq. (10), the slope of the regression line through the data points (*V*_*r*_, *V*_*d*_) gives an estimate of (1 − *α*_1_)(1 − *w*), from which *α*_1_ can be inferred.

### Master equation for the dynamics of gene product number

Let *p*_*k,n*_ denote the probability of observing microstate (*k, n*), where *k* is the cell cycle stage and *n* is the copy number of the gene product. Then the evolution of copy number dynamics is governed by the master equation

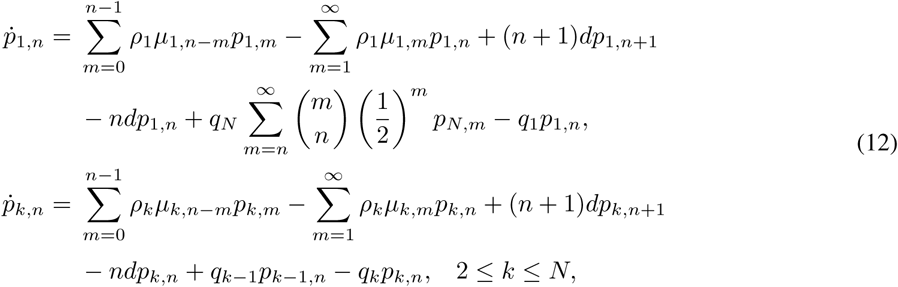

where *µ*_*k,n*_ = [*B*_*k*_/(*B*_*k*_ + 1)]^*n*^[1/(*B*_*k*_ + 1)] is the burst size distribution at stage *k*, the first two terms on the right-hand side describe bursty production, the middle two describe degradation, and the last two describe cell cycle progression or binomial partitioning of gene product molecules at cell division. The master equation for the concentration dynamics can also be derived in the large molecule number limit (Supplementary Note 2).

### Lineage distributions of gene product number

We consider the distribution of copy numbers for lineage measurements. Note that at each cell cycle stage, the stochastic dynamics of copy number is exactly the classical discrete bursty model proposed in [69], whose steady-state is the negative binomial distribution. Therefore, it is natural to approximate the conditional distribution of copy number at each stage by the negative binomial distribution

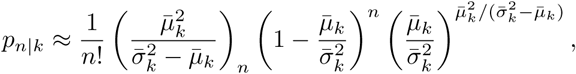

where 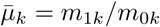 and 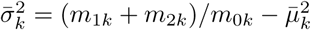 are the conditional mean and variance of copy number at stage *k*, respectively, and (*x*)_*n*_ = *x*(*x* + 1) *…* (*x* + *n* − 1) denotes the Pochhammer symbol. Hence the overall distribution of copy number is given by the following mixture of *N* negative binomial distributions:

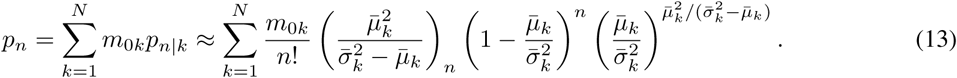

### Lineage distributions of gene product concentration

We consider the distribution of concentration for lineage measurements. Note that at each cell cycle stage, the stochastic dynamics of concentration is exactly the classical continuous bursty model proposed in [50], whose steady-state is the gamma distribution. Therefore, it is natural to approximate the conditional distribution of concentration at each stage by the gamma distribution

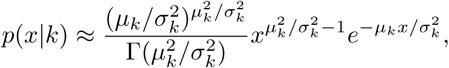

where *μ*_*k*_ = *m*_1*k*_/(*m*_0*k*_*v*_*k*_) and 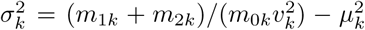 are the conditional mean and variance of concentration at stage *k*, respectively. Hence the overall distribution of concentration is given by the following mixture of *N* gamma distributions:

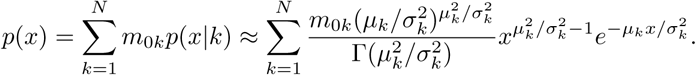

### Population distributions of gene product number and concentration

Up till now and in the main text, we have calculated gene product statistics for lineage measurements. Here we perform calculations for population measurements following the approach described in [14, 18]. Let *C*_*k*_(*t*) be the mean number of cells at stage *k* in the population at time *t*. Then all *C*_*k*_(*t*), 1 ≤ *r* ≤ *N* satisfy the following set of differential equations:

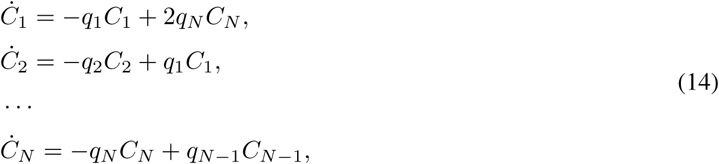

where the term 2*q*_*N*_ *C*_*N*_ in the first equation is due to the fact that the cell divides when the system transitions from stage *N* to stage 1. We then make the ansatz

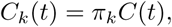

where *C*(*t*) is the total number of cells in the population and *π*_*k*_ is the proportion of cells at stage *k*. Then Eq. (14) can be rewritten as

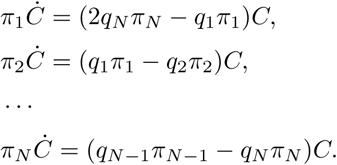

It then follows from the above equation that

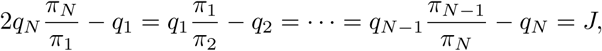

where *J* = *Ċ /C*. This clearly shows that

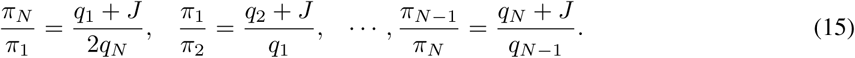

Multiplying all the above equations shows that *J* is the solution to the equation

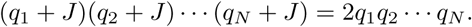

It thus follows from Eq. (15) that

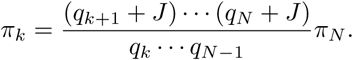

Since all *π*_*k*_ add up to 1, we finally obtain

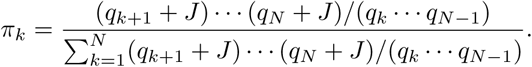

Now we have obtained the proportion of cells at each stage. Therefore, the population distribution of copy number is given by the following mixture of *N* negative binomial distributions:

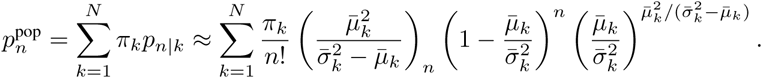

Similarly, the population distribution of concentration is given by the following mixture of *N* gamma distributions:

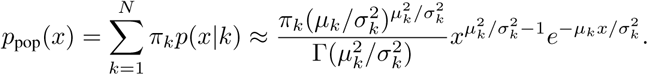

## Acknowledgments

C. J. acknowledges support from the NSAF grant in National Natural Science Foundation of China (No. U1930402). A. S. is supported by the National Institute of Health Grant 1R01GM126557. R. G. acknowledges support from the Leverhulme Trust (RPG-2020-327).

## Author contributions

R. G. conceived the original idea and directed the research. C. J. performed the theoretical derivations and numerical simulations. C. J, A. S, and R. G interpreted the theoretical results. C. J and R. G jointly wrote the manuscript with input from A. S.

## Data and materials availability

The MATLAB codes of stochastic simulations of the full and mean-field models can be found on GitHub via the link https://github.com/chenjiacsrc/Concentration-homeostasis. The experimental data used are from the papers [53, 57, 70].

## References

[1] Peccoud, J. & Ycart, B. Markovian modeling of gene-product synthesis. Theor. Popul. Biol. 48, 222–234 (1995).

[2] Shahrezaei, V. & Swain, P. S. Analytical distributions for stochastic gene expression. Proc. Natl. Acad. Sci. USA 105, 17256–17261 (2008).

[3] Smith, S. & Grima, R. Single-cell variability in multicellular organisms. Nat. Commun. 9, 1–8 (2018).

[4] Zenklusen, D., Larson, D. R. & Singer, R. H. Single-RNA counting reveals alternative modes of gene expression in yeast. Nat. Struct. Mol. Biol. 15, 1263–1271 (2008).

[5] Larsson, A. J. et al. Genomic encoding of transcriptional burst kinetics. Nature 565, 251–254 (2019).

[6] Taniguchi, Y. et al. Quantifying E. coli proteome and transcriptome with single-molecule sensitivity in single cells. Science 329, 533–538 (2010).

[7] Ham, L., Brackston, R. D. & Stumpf, M. P. Extrinsic noise and heavy-tailed laws in gene expression. Phys. Rev. Lett. 124, 108101 (2020).

[8] Kumar, N., Platini, T. & Kulkarni, R. V. Exact distributions for stochastic gene expression models with bursting and feedback. Phys. Rev. Lett. 113, 268105 (2014).

[9] Singh, A. Negative feedback through mRNA provides the best control of gene-expression noise. IEEE T. Nanobiosci. 10, 194–200 (2011).

[10] Jia, C. & Grima, R. Small protein number effects in stochastic models of autoregulated bursty gene expression. J. Chem. Phys. 152, 084115 (2020).

[11] To, T.-L. & Maheshri, N. Noise can induce bimodality in positive transcriptional feedback loops without bistability. Science 327, 1142–1145 (2010).

[12] Thomas, P., Popović, N. & Grima, R. Phenotypic switching in gene regulatory networks. Proc. Natl. Acad. Sci. USA 111, 6994–6999 (2014).

[13] Tanouchi, Y. et al. A noisy linear map underlies oscillations in cell size and gene expression in bacteria. Nature 523, 357–360 (2015).

[14] Beentjes, C. H., Perez-Carrasco, R. & Grima, R. Exact solution of stochastic gene expression models with bursting, cell cycle and replication dynamics. Phys. Rev. E 101, 032403 (2020).

[15] Johnston, I. G. & Jones, N. S. Closed-form stochastic solutions for non-equilibrium dynamics and inheritance of cellular components over many cell divisions. Proc. R. Soc. A 471, 20150050 (2015).

[16] Antunes, D. & Singh, A. Quantifying gene expression variability arising from randomness in cell division times. J. Math. Biol. 71, 437–463 (2015).

[17] Soltani, M., Vargas-Garcia, C. A., Antunes, D. & Singh, A. Intercellular variability in protein levels from stochastic expression and noisy cell cycle processes. PLoS Comput. Biol. 12, e1004972 (2016).

[18] Perez-Carrasco, R., Beentjes, C. & Grima, R. Effects of cell cycle variability on lineage and population measurements of messenger RNA abundance. J. R. Soc. Interface 17, 20200360 (2020).

[19] Dessalles, R., Fromion, V. & Robert, P. Models of protein production along the cell cycle: An investigation of possible sources of noise. PLoS one 15, e0226016 (2020).

[20] Jia, C. & Grima, R. Frequency domain analysis of fluctuations of mRNA and protein copy numbers within a cell lineage: theory and experimental validation. Phys. Rev. X 11, 021032 (2021).

[21] Jia, C., Singh, A. & Grima, R. Cell size distribution of lineage data: analytic results and parameter inference. iScience 24, 102220 (2021).

[22] Wang, M., Zhang, J., Xu, H. & Golding, I. Measuring transcription at a single gene copy reveals hidden drivers of bacterial individuality. Nat. Microbiol. 4, 2118–2127 (2019).

[23] Padovan-Merhar, O. et al. Single mammalian cells compensate for differences in cellular Volume and DNA copy number through independent global transcriptional mechanisms. Mol. Cell 58, 339–352 (2015).

[24] Sun, X.-M. et al. Size-dependent increase in RNA Polymerase II initiation rates mediates gene expression scaling with cell size. Curr. Biol. 30, 1217–1230 (2020).

[25] Kempe, H., Schwabe, A., Crémazy, F., Verschure, P. J. & Bruggeman, F. J. The volumes and transcript counts of single cells reveal concentration homeostasis and capture biological noise. Mol. Biol. Cell 26, 797–804 (2015).

[26] Facchetti, G., Chang, F. & Howard, M. Controlling cell size through sizer mechanisms. Curr. Opin. Syst. Biol. 5, 86–92 (2017).

[27] Ghusinga, K. R., Vargas-Garcia, C. A. & Singh, A. A mechanistic stochastic framework for regulating bacterial cell division. Sci. Rep. 6, 1–9 (2016).

[28] Sekar, K. et al. Synthesis and degradation of FtsZ quantitatively predict the first cell division in starved bacteria. Mol. Syst. Biol. 14, e8623 (2018).

[29] Si, F. et al. Mechanistic origin of cell-size control and homeostasis in bacteria. Curr. Biol. 29, 1760–1770 (2019).

[30] Pan, K. Z., Saunders, T. E., Flor-Parra, I., Howard, M. & Chang, F. Cortical regulation of cell size by a sizer cdr2p. Elife 3, e02040 (2014).

[31] Keifenheim, D. et al. Size-dependent expression of the mitotic activator Cdc25 suggests a mechanism of size control in fission yeast. Curr. Biol. 27, 1491–1497 (2017).

[32] Facchetti, G., Knapp, B., Flor-Parra, I., Chang, F. & Howard, M. Reprogramming Cdr2-dependent geometry-based cell size control in fission yeast. Curr. Biol. 29, 350–358 (2019).

[33] Patterson, J. O., Rees, P. & Nurse, P. Noisy cell-size-correlated expression of cyclin b drives probabilistic cell-size homeostasis in fission yeast. Curr. Biol. 29, 1379–1386 (2019).

[34] Wang, P. et al. Robust growth of Escherichia coli. Curr. Biol. 20, 1099–1103 (2010).

[35] Soifer, I., Robert, L. & Amir, A. Single-cell analysis of growth in budding yeast and bacteria reveals a common size regulation strategy. Curr. Biol. 26, 356–361 (2016).

[36] Cermak, N. et al. High-throughput measurement of single-cell growth rates using serial microfluidic mass sensor arrays. Nat. Biotechnol. 34, 1052–1059 (2016).

[37] Priestman, M., Thomas, P., Robertson, B. D. & Shahrezaei, V. Mycobacteria modify their cell size control under sub-optimal carbon sources. Front. Cell Dev. Biol. 5, 64 (2017).

[38] Eun, Y.-J. et al. Archaeal cells share common size control with bacteria despite noisier growth and division. Nat. Microbiol. 3, 148–154 (2018).

[39] Jia, C., Yin, G. G., Zhang, M. Q. et al. Single-cell stochastic gene expression kinetics with coupled positive-plus-negative feedback. Phys. Rev. E 100, 052406 (2019).

[40] Chao, H. X. et al. Evidence that the human cell cycle is a series of uncoupled, memoryless phases. Mol. Syst. Biol. 15 (2019).

[41] Wood, E. & Nurse, P. Pom1 and cell size homeostasis in fission yeast. Cell Cycle 12, 3417–3425 (2013).

[42] Schmoller, K. M. The phenomenology of cell size control. Curr. Opin. Cell Biol. 49, 53–58 (2017).

[43] Chandler-Brown, D., Schmoller, K. M., Winetraub, Y. & Skotheim, J. M. The adder phenomenon emerges from independent control of pre-and post-start phases of the budding yeast cell cycle. Curr. Biol. 27, 2774–2783 (2017).

[44] Cadart, C. et al. Size control in mammalian cells involves modulation of both growth rate and cell cycle duration. Nat. Commun. 9, 1–15 (2018).

[45] Nieto, C., Arias-Castro, J., Sánchez, C., Vargas-García, C. & Pedraza, J. M. Unification of cell division control strategies through continuous rate models. Phys. Rev. E 101, 022401 (2020).

[46] Skinner, S. O. et al. Single-cell analysis of transcription kinetics across the cell cycle. Elife 5, e12175 (2016).

[47] Dowling, M. R. et al. Stretched cell cycle model for proliferating lymphocytes. Proc. Natl. Acad. Sci. USA 111, 6377–6382 (2014).

[48] Kadanoff, L. P. Statistical physics: statics, dynamics and renormalization (World Scientific Publishing Company, 2000).

[49] Yin, G. & Zhu, C. Hybrid switching diffusions: properties and applications, vol. 63 (Springer, New York, 2010).

[50] Friedman, N., Cai, L. & Xie, X. S. Linking stochastic dynamics to population distribution: an analytical framework of gene expression. Phys. Rev. Lett. 97, 168302 (2006).

[51] Lin, J. & Amir, A. Homeostasis of protein and mRNA concentrations in growing cells. Nat. Commun. 9, 1–11 (2018).

[52] Jia, C., Singh, A. & Grima, R. Characterizing non-exponential growth and bimodal cell size distributions in Schizosaccharomyces pombe: an analytical approach. bioRxiv (2021).

[53] Claude, K.-L. et al. Transcription coordinates histone amounts and genome content. Nat. Commun. 12, 1–17 (2021).

[54] Wang, Y. et al. Precision and functional specificity in mRNA decay. Proc. Natl. Acad. Sci. USA 99, 5860–5865 (2002).

[55] Khmelinskii, A. et al. Tandem fluorescent protein timers for in vivo analysis of protein dynamics. Nat. Biotechnol. 30, 708 (2012).

[56] Christiano, R., Nagaraj, N., Fröhlich, F. & Walther, T. C. Global proteome turnover analyses of the yeasts S. cerevisiae and S. pombe. Cell Rep. 9, 1959–1965 (2014).

[57] Nakaoka, H. & Wakamoto, Y. Aging, mortality, and the fast growth trade-off of Schizosaccharomyces pombe. PLoS Biol. 15, e2001109 (2017).

[58] Cao, Z. & Grima, R. Analytical distributions for detailed models of stochastic gene expression in eukaryotic cells. Proc. Natl. Acad. Sci. USA 117, 4682–4692 (2020).

[59] Thomas, P. & Shahrezaei, V. Coordination of gene expression noise with cell size: extrinsic noise versus agent-based models of growing cell populations. J. R. Soc. Interface 18, 20210274 (2021).

[60] Van Kampen, N. G. Stochastic processes in physics and chemistry, vol. 1 (Elsevier, 1992).

[61] Grima, R. Linear-noise approximation and the chemical master equation agree up to second-order moments for a class of chemical systems. Phys. Rev. E 92, 042124 (2015).

[62] Taheri-Araghi, S. et al. Cell-size control and homeostasis in bacteria. Curr. Biol. 25, 385–391 (2015).

[63] Liu, X., Oh, S., Peshkin, L. & Kirschner, M. W. Computationally enhanced quantitative phase microscopy reveals autonomous oscillations in mammalian cell growth. Proc. Natl. Acad. Sci. USA 117, 27388–27399 (2020).

[64] Tian, T. & Burrage, K. Stochastic models for regulatory networks of the genetic toggle switch. Proc. Natl. Acad. Sci. USA 103, 8372–8377 (2006).

[65] Singh, A. & Hespanha, J. P. Optimal feedback strength for noise suppression in autoregulatory gene networks. Biophys. J. 96, 4013–4023 (2009).

[66] Chalancon, G. et al. Interplay between gene expression noise and regulatory network architecture. Trends Genet. 28, 221–232 (2012).

[67] Holehouse, J., Cao, Z. & Grima, R. Stochastic modeling of autoregulatory genetic feedback loops: A review and comparative study. Biophys. J. 118, 1517–1525 (2020).

[68] Cao, Z. & Grima, R. Linear mapping approximation of gene regulatory networks with stochastic dynamics. Nat. Commun. 9, 3305 (2018).

[69] Paulsson, J. & Ehrenberg, M. Random signal fluctuations can reduce random fluctuations in regulated components of chemical regulatory networks. Phys. Rev. Lett. 84, 5447 (2000).

[70] Tanouchi, Y. et al. Long-term growth data of Escherichia coli at a single-cell level. Sci. Data 4, 1–5 (2017).

