## Supplementary material for "Concentration fluctuations due to size-dependent gene expression and cell-size control mechanisms"

**4 Technical details for Fig. 3 15**

**5 Technical details for Fig. 4 15**

**6 Technical details for Fig. 5 16**

#### Note 1. Derivation of the moment expressions for the mean-field model

##### 1.1 Moments of gene product number

For each cell cycle stage $k$, we introduce the generating function

$$F_k(z) = \sum_{n=0}^{\infty} p_{k,n} z^n.$$

Then the master equation of the mean-field model (see Eq. (9) in the main text) can be converted into the following system of partial differential equations:

$$\begin{aligned}
\partial_t F_1(z) &= \rho_1[H_1(z) - H_1(1)]F_1(z) \\
&\quad + d(1-z)\partial_z F_1(z) + q_N F_N\left(\frac{z+1}{2}\right) - q_1 F_1(z), \\
\partial_t F_k(z) &= \rho_k[H_k(z) - H_k(1)]F_k(z) \\
&\quad + d(1-z)\partial_z F_k(z) + q_{k-1}F_{k-1}(z) - q_k F_k(z), \quad 2 \le k \le N.
\end{aligned} \tag{1}$$

where

$$H_k(z) = \sum_{n=1}^{\infty} \mu_{k,n} z^n = \frac{B_k z}{(B_k+1)[B_k(1-z)+1]},$$

is the generating function of the typical burst size distribution $\mu_k = (\mu_{k,n})$ at stage $k$. Recall that for each $l \geq 0$, the $l$th unnormalized factorial moment of gene product number at stage $k$ is defined as

$$m_{lk} := \sum_{n=0}^{\infty} n(n-1)\cdots(n-l+1)p_{k,n} = F_k^{(l)}(1).$$

In particular, $m_{0k} = \sum_{n=0}^{\infty} p_{k,n}$ is the probability of being at stage $k$. To proceed, let $m_l = (m_{lk})$ be the row vector whose components are the $l$th factorial moments at all stages. Taking the $k$th derivative on both sides of Eq. (1), we obtain

$$\dot{m}_l = m_l W_{ll} + \sum_{j=0}^{l-1} m_j W_{jl}, \tag{2}$$

where $W_{ll}$ and $W_{jl}$ are matrices defined as

$$W_{ll} = \begin{pmatrix} -q_1 & q_1 & & & \\ & -q_2 & q_2 & & \\ & & \ddots & \ddots & \\ & & & -q_{N-1} & q_{N-1} \\ q_N/2^l & & & & -q_N \end{pmatrix} - ldI, \quad l \geq 0, \tag{3}$$

with $I$ being the identity matrix, and

$$W_{jl} = C_{l,j}\mathrm{diag}(H_1^{(l-j)}(1)\rho_1, \cdots, H_N^{(l-j)}(1)\rho_N), \quad j \leq l-1.$$

At the steady state, we have $m_0 W_{00} = 0$ and its solution is given by

$$m_{0k} = \frac{1/q_k}{\sum_{i=1}^N 1/q_i}.$$

Then the higher-order factorial moments of gene product number can be computed recursively as

$$m_l = -\sum_{j=0}^{l-1} m_j W_{jl} W_{ll}^{-1}, \quad l \geq 1.$$

In particular, the first and second factorial moments are given by

$$\begin{aligned} m_1 &= -m_0 W_{01} W_{11}^{-1}, \\ m_2 &= -(m_1 W_{12} + m_0 W_{02}) W_{22}^{-1}. \end{aligned} \tag{4}$$

Therefore, the mean $\langle n \rangle$ and variance $\sigma_n^2$ of gene product number are given by

$$\langle n \rangle = m_1 \mathbb{1} = \sum_{k=1}^N m_{1k}, \quad \sigma_n^2 = (m_1+m_2)\mathbb{1} - (m_1\mathbb{1})^2 = \sum_{k=1}^N (m_{1k}+m_{2k}) - \langle n \rangle^2, \tag{5}$$

where $\mathbb{1} = (1, \cdots, 1)^T$ is the column vector whose components are all 1. It is easy to check that

$$W_{01} = SH, \quad W_{12} = 2SH, \quad W_{02} = 2SH^2,$$

where $S = \mathrm{diag}(\rho_1, \cdots, \rho_N)$ is the diagonal matrix whose diagonal entries are the burst frequencies at all stages and $H = \mathrm{diag}(B_1, \cdots, B_N)$ is the diagonal matrix whose diagonal entries are the typical mean burst sizes at all stages, which may or may not scale with cell size.

#### 1.2 Moments of gene product concentration

Let $\mu_{lk}$ be the unnormalized $l$th moment of gene product concentration at stage $k$ and let $\mu_l = (\mu_{lk})$ be the row vector whose components are the $l$th moments of concentration at all stages. It is easy to see that

$$\mu_1 = m_1 V^{-1}, \quad \mu_2 = (m_1 + m_2)V^{-2},$$

where $V = \mathrm{diag}(v_1, \cdots, v_N)$ is the diagonal matrix whose diagonal entries are the typical cell sizes at all stages. The components of $\mu_1$ and $\mu_2$ can be written as

$$\mu_{1k} = \frac{m_{1k}}{v_k}, \quad \mu_{2k} = \frac{m_{1k} + m_{2k}}{v_k^2}.$$

At the steady state, it follows from Eq. (4) that

$$\begin{aligned} \mu_1 &= -m_0 W_{01} W_{11}^{-1} V^{-1}, \\ \mu_2 &= -[m_0 W_{01} W_{11}^{-1} + (m_1 W_{12} + m_0 W_{02}) W_{22}^{-1}] V^{-2}. \end{aligned} \tag{6}$$

Therefore, the mean $\langle c \rangle$ and variance $\sigma_c^2$ of gene product concentration are given by

$$\langle c \rangle = \mu_1 \mathbb{1} = \sum_{k=1}^{N} \frac{m_{1k}}{v_k}, \quad \sigma_c^2 = \mu_2 \mathbb{1} - (\mu_1 \mathbb{1})^2 = \sum_{k=1}^{N} \frac{m_{1k} + m_{2k}}{v_k^2} - \langle c \rangle^2. \tag{7}$$

#### 1.3 Noise in gene product number and concentration

Once we have obtained the expressions of first and second moments, we can provide a detailed analysis of gene expression noise. Recall that noise $\phi_n$ in gene product number and noise $\phi_c$ in gene product concentration are defined as

$$\phi_n = \frac{\sigma_n^2}{\langle n \rangle^2}, \quad \phi_c = \frac{\sigma_c^2}{\langle c \rangle^2}.$$

From Eq. (5), noise $\phi_n$ in copy number can be decomposed as

$$\phi_n = \phi_n^{\text{intrisic}} + \phi_n^{\text{extrisic}}, \tag{8}$$

where

$$\phi_n^{\text{intrisic}} = \frac{1}{\langle n \rangle} \left[ 1 + \frac{m_0 W_{02} W_{22}^{-1} \mathbb{1}}{m_0 W_{01} W_{11}^{-1} \mathbb{1}} \right],$$

is the intrinsic noise and

$$\phi_n^{\text{extrisic}} = \frac{m_0 W_{01} W_{11}^{-1} W_{12} W_{22}^{-1} \mathbb{1}}{(m_0 W_{01} W_{11}^{-1} \mathbb{1})^2} - 1,$$

is the extrinsic noise. It is easy to see that both the extrinsic noise and the term in the bracket of the intrinsic noise are independent of the burst frequency $\rho$. Moreover, the intrinsic noise is the sum of two terms: the term 1 in the bracket represents Poissonian noise and the other term, which is proportional to the mean burst size $B$, represents noise due to bursting.

Similarly, it follows from Eq. (7) that noise $\phi_c$ in concentration can be decomposed as

$$\phi_c = \phi_c^{\text{intrisic}} + \phi_c^{\text{extrisic}}, \tag{9}$$

where

$$\phi_c^{\text{intrisic}} = \frac{1}{\langle c \rangle} \left[ \frac{m_0 W_{01} W_{11}^{-1} V^{-2} \mathbb{1}}{m_0 W_{01} W_{11}^{-1} V^{-1} \mathbb{1}} + \frac{m_0 W_{02} W_{22}^{-1} V^{-2} \mathbb{1}}{m_0 W_{01} W_{11}^{-1} V^{-1} \mathbb{1}} \right],$$

is the intrinsic noise and

$$\phi_c^{\text{extrisic}} = \frac{m_0 W_{01} W_{11}^{-1} W_{12} W_{22}^{-1} V^{-2} \mathbb{1}}{(m_0 W_{01} W_{11}^{-1} V^{-1} \mathbb{1})^2} - 1,$$

is the extrinsic noise. Similarly, it is easy to see that both the extrinsic noise and the term in the bracket of the intrinsic noise are independent of the burst frequency $\rho$. Moreover, the intrinsic noise is the sum of two terms: the first term in the bracket represents noise from individual births and deaths of the gene product, and the other term, which is proportional to the mean burst size $B$, represents noise due to bursting.

##### 1.4 Perfect concentration homeostasis

Concentration homeostasis is perfect when the mean concentration at each cell cycle stage is a constant. Here we shall prove that perfect homeostasis is achieved when $\beta = \kappa = 1$.

Before studying perfect homeostasis, we note that the cell cycle duration $T$ for the mean-field model is the independent sum of $N$ exponentially distributed random variables with rates $q_1, \cdots, q_N$, respectively. Practically, this distribution is well approximated by an Erlang distribution [1]. The mean and variance of the cell cycle duration can be easily computed as

$$\langle T \rangle = \frac{1}{q_1} + \cdots + \frac{1}{q_N}, \quad \sigma_T^2 = \frac{1}{q_1^2} + \cdots + \frac{1}{q_N^2}.$$

If we use an Erlang distribution with shape parameter $\bar{N}$ and rate $\bar{a}$ to approximate this distribution, then $\bar{N}$ and $\bar{a}$ should satisfy

$$\frac{\bar{N}}{\bar{a}} = \langle T \rangle, \quad \frac{\bar{N}}{\bar{a}^2} = \sigma_T^2.$$

Thus the two parameters can be determined as

$$\bar{N} = \frac{\langle T \rangle^2}{\sigma_T^2}, \quad \bar{a} = \frac{\langle T \rangle}{\sigma_T^2} = \bar{N} f.$$

Note that for the timer strategy, i.e. $\alpha_0, \alpha_1 \to 0$, the transition rate between stages is a constant and thus the cell cycle duration is exactly Erlang distributed. The above discussion suggests that the mean-field model for an arbitrary size control strategy can be well approximated by a mean-field model for the timer strategy with $\bar{N}$ cell cycle stages and transition rate $\bar{a} = \bar{N} f$ between stages. The parameter $N_0$ for the effective timer model can then be determined as $\bar{N}_0 = w\bar{N}$, where $w = \log_2(v_{N_0+1}/v_1)$ is the proportion of cell cycle before replication. Therefore, in order to investigate an arbitrary size control strategy, we only need to investigate the timer strategy first and then replace the parameters $N$ and $N_0$ in the effective timer model by $\bar{N}$ and $\bar{N}_0$, respectively.

We next examine perfect concentration homeostasis for the timer strategy. In this case, the transition rate between stages is a constant and we denote it by $\bar{a} = Nf$. Since both $W_{00}$ and $W_{11}$ are circular matrices, their eigenvalues can be computed explicitly. The eigenvalues of $W_{00}$ are given by

$$\lambda_k = -\bar{a} + \bar{a}\omega_k, \quad 1 \le k \le N, \tag{10}$$

and the eigenvalues of $W_{11}$ are given by

$$\lambda_{N+k} = -d - \bar{a} + 2^{-1/N}\bar{a}\omega_k, \quad 1 \le k \le N, \tag{11}$$

where $\omega_k = e^{2(k-1)\pi i/N}$ are all the $N$th roots of unity. Since $W_{00}$ is a normal matrix, it is easy to check that there exists a complex orthogonal matrix

$$R = \frac{1}{\sqrt{N}} \begin{pmatrix} 1 & 1 & \cdots & 1 \\ \omega_1 & \omega_2 & \cdots & \omega_N \\ \cdots & \cdots & \cdots & \cdots \\ \omega_1^{N-1} & \omega_2^{N-1} & \cdots & \omega_N^{N-1} \end{pmatrix},$$

such that $W_{00}$ is diagonalized, i.e.

$$W_{00} = RD_{00}\bar{R}', \tag{12}$$

where $D_{00} = \mathrm{diag}(\lambda_1, \cdots, \lambda_N)$ and $\bar{R}'$ denotes the conjugate transpose of $R$. Similarly, $W_{11}$ can be diagonalized as

$$W_{11} = MRD_{11}\bar{R}'M^{-1}, \tag{13}$$

where $D_{11} = \mathrm{diag}(\lambda_{N+1}, \cdots, \lambda_{2N})$ and $M$ is the diagonal matrix given by

$$M = \mathrm{diag}(1, 2^{-1/N}, \cdots, 2^{-(N-1)/N}).$$

It is clear that the matrices $V$ and $M$ are related by

$$V = v_1 M^{-1}. \tag{14}$$

For the timer strategy, the vector $m_0$ can be computed explicitly as

$$m_0 = \frac{1}{N}\mathbb{1}'. \tag{15}$$

Combining Eqs. (13), (14), and (15), the vector $\mu_1$ can be rewritten as

$$\mu_1 = -m_0 W_{01} W_{11}^{-1} V^{-1} = -\frac{1}{Nv_1}\mathbb{1}' W_{01} MRD_{11}^{-1}\bar{R}'.$$

This can be written in components as

$$\mu_{1k} = -\frac{1}{Nv_1}\sum_{j,l=1}^{N} \frac{[W_{01}M]_{jj} R_{jl}\bar{R}_{kl}}{\lambda_{N+l}} = \frac{1}{Nv_1}\sum_{j,l=1}^{N} \frac{[W_{01}M]_{jj} R_{jl}\bar{R}_{kl}}{d + \bar{a} - 2^{-1/N}\bar{a}\omega_l},$$

where $A_{jl}$ denotes the $(j,l)$-th entry of the matrix $A$. It is easy to check that

$$W_{01} = SH = \rho B v_1^{\beta} \mathrm{diag}(1, \cdots, 2^{\beta(N_0-1)/N}, \kappa 2^{\beta N_0/N}, \cdots, \kappa 2^{\beta(N-1)/N}).$$

When $\beta = \kappa = 1$, we have $W_{01} = \rho B v_1 M^{-1}$ and thus $W_{01}M = \rho B v_1 I$ is a constant multiple of the identity matrix. This clearly shows that

$$\mu_{1k} = \frac{\rho B}{N}\sum_{j,l=1}^{N} \frac{R_{jl}\bar{R}_{kl}}{d + \bar{a} - 2^{-1/N}\bar{a}\omega_l}. \tag{16}$$

We next make a crucial observation that

$$\sum_{j=1}^{N} R_{jl} = \frac{1}{\sqrt{N}}\sum_{j=1}^{N} \omega_l^{j-1} = \sqrt{N}\delta_{l1}.$$

Inserting this equation into Eq. (16) yields

$$\mu_{1k} = \frac{\rho B}{\sqrt{N}} \frac{\bar{R}_{k1}}{d + \bar{a} - 2^{-1/N}\bar{a}\omega_1} = \frac{\rho B}{N(d + \bar{a} - 2^{-1/N}\bar{a})}.$$

Note that when $N \gg 1$, we have

$$d + \bar{a} - 2^{-1/N}\bar{a} = d + (1 - e^{-\log(2)/N})Nf \approx d + (\log 2)f = d_{\text{eff}}.$$

Therefore, the unnormalized mean concentration $\mu_{1k}$ at stage $k$ is given by

$$\mu_{1k} = \frac{\rho B}{N d_{\text{eff}}},$$

and the normalized mean concentration $\mu_{1k}$ at stage $k$ is given by

$$\mu_k = \frac{\mu_{1k}}{m_{0k}} = \frac{\rho B}{d_{\text{eff}}},$$

which is independent of stage $k$. As a result, we have proved that perfect concentration homeostasis is achieved when $\beta = \kappa = 1$.

#### Note 2. Mean-field model of the concentration dynamics

Note that the dynamics of gene product number at each cell cycle stage is the classical discrete bursty model proposed in [2]. It is interesting to understand the dynamics of gene product concentration at each stage. To the end, we make the approximation of large molecule number. Recall that the burst size distribution $\mu = (\mu_n)$ is given by $\mu_n = p_B^n(1 - p_B)$, where $p_B = B'/(B' + 1)$ with $B' = BV(t)^\beta$ being the mean burst size. When the mean burst size $B'$ is large, we have

$$-\log p_B = -\log(1 - 1/(B' + 1)) \approx 1/(B' + 1) = 1 - p_B$$

and thus the burst size distribution is given by

$$\mu_n = p_B^n(1 - p_B) = e^{n \log p_B}(1 - p_B) \approx e^{-(1-p_B)n}(1 - p_B),$$

which is approximately an exponential distribution with mean $1/(1 - p_B) = (B' + 1) \approx B'$. As a result, the synthesis of the gene product at stage $k$ can be described by a compound Poisson process with exponentially distributed interarrival times with rate $\rho_k$ and exponentially distributed burst sizes with mean $B'$. By the scaling property of the exponential distribution, an exponentially distributed random variable with mean $B' = BV(t)^\beta$ can be viewed as an exponentially distributed random variable with mean $B$ multiplied by $V(t)^\beta$. Therefore, in the large burst size limit, the stochastic gene expression dynamics at stage $k$ can be approximated by the stochastic differential equation (SDE) [3]

$$\dot{n}(t) = V(t)^\beta \dot{s}_k(t) - dn(t), \tag{17}$$

where $n(t)$ denotes the gene product number at time $t$ and $s_k(t)$ denotes a compound Poisson process with arrival rate $\rho_k$ and an exponentially distributed jump distribution with mean $B$.

Let $c(t) = n(t)/V(t)$ denote the gene product concentration at time $t$. Then Eq. (17) can be written as

$$V(t)\dot{c}(t) + gV(t)c(t) = V(t)^\beta \dot{s}_k(t) - dV(t)c(t),$$

where we have used the fact that $\dot{V}(t) = gV(t)$ since cell size grows exponentially with rate $g$. Thus the dynamics of gene product concentration at stage $k$ is governed by the SDE

$$\dot{c}(t) = V(t)^{\beta-1}\dot{s}_k(t) - (d+g)c(t) = V(t)^{\beta-1}\dot{s}_k(t) - d_{\text{eff}}c(t),$$

where $d_{\text{eff}} = d+g \approx d+\log(2)f$ is the effective decay rate of the gene product due to active degradation and dilution at cell division. Under the mean-field approximation, we have $V(t) \approx v_k$ and thus the concentration dynamics at stage $k$ can be described by the continuous bursty model

$$\dot{c}(t) = v_k^{\beta-1}\dot{s}_k(t) - d_{\text{eff}}c(t). \tag{18}$$

The remaining question is whether cell division affects the concentration dynamics. Actually, in the large molecule number limit, the binomial partitioning of molecule number reduces to deterministic partitioning. This is a direct consequence of the law of large numbers since a binomial random variable can be viewed as the i.i.d. sum of Bernoulli random variables. At the moment of cell division, both the molecule number and the cell volume undergo deterministic symmetric partitioning, and thus their ratio remains invariant. This shows that in the large molecule number limit, cell division has no effect on concentration fluctuations.

Now we can construct a model describing concentration fluctuations across the cell cycle. At each stage $k$, the concentration dynamics is governed by Eq. (18) and the system can hop from stage $k$ to the next with rate $q_k$. The stochastic dynamics of this system are described by a hybrid Markovian model whose Kolmogorov forward equation is given by [3, 4]

$$\begin{aligned}\partial_t p_1(x) &= d_{\text{eff}}\partial_x\big(xp(x)\big) + \rho_1\int_0^x w_1(x-y)p(y)dy - \rho_1 p(x)\\ &\quad + q_N p_N(x) - q_1 p_1(x),\\ \partial_t p_k(x) &= d_{\text{eff}}\partial_x\big(xp(x)\big) + \rho_k\int_0^x w_k(x-y)p(y)dy - \rho_k p(x)\\ &\quad + q_{k-1}p_{k-1}(x) - q_k p_k(x),\quad 2\leq k\leq N,\end{aligned} \tag{19}$$

where $p_k(x)$ is the probability density of concentration when the cell is at stage $k$ and

$$w_k(x) = \frac{1}{Bv_k^{\beta-1}}e^{-x/(Bv_k^{\beta-1})}.$$

is the burst size distribution at stage $k$. A special case occurs when the synthesis is balanced ($\beta = 1$) and dosage compensation is perfect ($\kappa = 1$). In this case, both the burst frequency $\rho_k = \rho$ and the burst size distribution $w_k(x) = (1/B)e^{-x/B}$ are independent of cell cycle stage $k$, and thus the concentration dynamics along the whole cell lineage is governed by

$$\partial_t p(x) = d_{\text{eff}}\partial_x\big(xp(x)\big) + \rho\int_0^x w(x-y)p(y)dy - \rho p(x), \tag{20}$$

where $p(x)$ is the probability density of concentration. This is exactly the classical continuous bursty model proposed by Friedman et al. [5].

In summary, in the large burst size limit, the concentration dynamics of our model reduces to the classical continuous gene expression model in the presence of balanced biosynthesis and perfect dosage compensation ($\beta = \kappa = 1$). In this case, the steady-state distribution of concentration can be derived from Eq. (20) and is given by [5]

$$p(x) = \frac{1}{B^{\rho/d_{\text{eff}}}\Gamma(\rho/d_{\text{eff}})}x^{\rho/d_{\text{eff}}-1}e^{-x/B},$$

which is a gamma distribution. In this case, the concentration mean $\rho B/d_{\text{eff}}$ and variance $\rho B^2/d_{\text{eff}}$ are both independent of cell cycle stage and thus are both independent of cell volume.

### Note 3. Derivation of the power spectra expressions for the mean-field model

#### 3.1 Power spectrum of gene product number

Let $r(t)$ denote the cell cycle stage and let $n(t)$ denote the gene product number in a single cell at time $t$. To proceed, let

$$\begin{aligned} m_{0k}(t) &= \sum_{n=0}^{\infty} p_{k,n} = \mathbb{P}(r(t)=k), \\ m_{1k}(t) &= \sum_{n=0}^{\infty} n p_{k,n} = \mathbb{E}n(t) I_{\{r(t)=k\}}, \\ m_{2k}(t) &= \sum_{n=0}^{\infty} n(n-1) p_{k,n} = \mathbb{E}n(t)(n(t)-1) I_{\{r(t)=k\}}, \end{aligned}$$

be the zeroth, first, and second factorial moments of gene product number at stage $k$, where $I_A$ denotes the indicator function of the set $A$. For convenience, let $m_k(t) = (m_{kr}(t))$ be the row vector whose components are the $k$th factorial moments at all stages. It then follows from Eq. (2) that $m_0(t)$, $m_1(t)$, and $m_2(t)$ satisfy the following differential equations:

$$\begin{aligned} \dot{m}_0(t) &= m_0(t) W_{00}, \\ \dot{m}_1(t) &= m_1(t) W_{11} + m_0(t) W_{01}, \\ \dot{m}_2(t) &= m_2(t) W_{22} + m_1(t) W_{12} + m_0(t) W_{02}. \end{aligned} \tag{21}$$

Since Eq. (21) is a set of linear differential equations, its time-dependent solution is given by

$$m_1(t) = m_1(0) e^{W_{11} t} + \int_0^t m_0(0) e^{W_{00} s} W_{01} e^{W_{11}(t-s)} \mathrm{d}s. \tag{22}$$

Given the initial cell cycle stage $r(0) = k$ and initial gene product number $n(0) = n$, it follows that

$$\mathbb{E}n(t) = m_1(t)\mathbb{1} = n e_k e^{W_{11} t}\mathbb{1} + \int_0^t e_k e^{W_{00} s} W_{01} e^{W_{11}(t-s)} \mathbb{1} \mathrm{d}s,$$

where $e_k$ denotes the row vector whose $k$th component is 1 and all other components are 0. This clearly shows that

$$\mathbb{E}[n(t)|r(0), n(0)] = n(0) e_{r(0)} e^{W_{11} t}\mathbb{1} + \int_0^t e_{r(0)} e^{W_{00} s} W_{01} e^{W_{11}(t-s)} \mathbb{1} \mathrm{d}s.$$

From now on, we assume that the system has reached the steady state. Then we have

$$\begin{aligned}
&\mathbb{E}n(0)n(t)\\
&= \sum_k \mathbb{E}n(0)I_{\{r(0)=k\}}\mathbb{E}[n(t)|r(0),n(0)]\\
&= \sum_k \mathbb{E}n(0)I_{\{r(0)=k\}}\left[n(0)e_{r(0)}e^{W_{11}t}\mathbb{1}+\int_0^t e_{r(0)}e^{W_{00}s}W_{01}e^{W_{11}(t-s)}\mathbb{1}\mathrm{d}s\right]\\
&= \sum_k \mathbb{E}n(0)^2I_{\{r(0)=k\}}e_ke^{W_{11}t}\mathbb{1}+\int_0^t \mathbb{E}n(0)I_{\{r(0)=k\}}e_ke^{W_{00}s}W_{01}e^{W_{11}(t-s)}\mathbb{1}\mathrm{d}s\\
&= \sum_k (m_{1k}+m_{2k})e_ke^{W_{11}t}\mathbb{1}+\int_0^t m_{1k}e_ke^{W_{00}s}W_{01}e^{W_{11}(t-s)}\mathbb{1}\mathrm{d}s\\
&= (m_1+m_2)e^{W_{11}t}\mathbb{1}+\int_0^t m_1e^{W_{00}s}W_{01}e^{W_{11}(t-s)}\mathbb{1}\mathrm{d}s,
\end{aligned}$$

where $m_1$ and $m_2$ are the steady-state values of moments given in Eq. (4). Since the autocorrelation function is defined as $R_n(t)=\mathbb{E}n(0)n(t)-\mathbb{E}n(0)\mathbb{E}n(t)$, we finally obtain an explicit expression of the autocorrelation function, which is given by

$$R_n(t)=(m_1+m_2)e^{W_{11}t}\mathbb{1}+\int_0^t m_1e^{W_{00}s}W_{01}e^{W_{11}(t-s)}\mathbb{1}\mathrm{d}s-(m_1\mathbb{1})^2.$$

##### 3.2 Power spectrum of gene product concentration

Let $c(t)$ denote the gene product concentration in a single cell at time $t$. To proceed, let

$$\begin{aligned}
\mu_{1k}(t) &= \mathbb{E}c(t)I_{\{r(t)=k\}} = m_{1k}(t)/v_k,\\
\mu_{2k}(t) &= \mathbb{E}c(t)^2I_{\{r(t)=k\}} = (m_{1k}(t)+m_{2k}(t))/v_k^2,
\end{aligned}$$

be the first and second moments of concentration at stage $k$. For convenience, let $\mu_k(t)=(\mu_{kr}(t))$ denote the row vector whose components are the $k$th moments of concentration at all stages. It is easy to see that

$$\mu_1(t)=m_1(t)V^{-1},\quad \mu_2(t)\approx[m_1(t)+m_2(t)]V^{-2}.$$

It then follows from Eq. (21) that

$$\mu_1(t)=m_1(0)e^{W_{11}t}V^{-1}+\int_0^t m_0(0)e^{W_{00}s}W_{01}e^{W_{11}(t-s)}V^{-1}\mathrm{d}s.$$

Given the initial cell cycle stage $r(0)=k$ and initial gene product number $n(0)=n$, it follows that

$$\mathbb{E}c(t)=\mu_1(t)\mathbb{1}=ne_ke^{W_{11}t}V^{-1}\mathbb{1}+\int_0^t e_ke^{W_{00}s}W_{01}e^{W_{11}(t-s)}V^{-1}\mathbb{1}\mathrm{d}s.$$

This clearly shows that

$$\mathbb{E}[c(t)|r(0),n(0)]\approx n(0)e_{r(0)}e^{W_{11}t}V^{-1}\mathbb{1}+\int_0^t e_{r(0)}e^{W_{00}s}W_{01}e^{W_{11}(t-s)}V^{-1}\mathbb{1}\mathrm{d}s.$$

From now on, we assume that the system has reached the steady state. Then we have

$$
\begin{aligned}
&\mathbb{E}c(0)c(t) \\
&= \sum_k \mathbb{E}c(0)I_{\{r(0)=k\}}\mathbb{E}[c(t)|r(0),n(0)] \\
&= \sum_k \mathbb{E}c(0)I_{\{r(0)=k\}}\left[n(0)e_{r(0)}e^{W_{11}t}V^{-1}\mathbb{1}+\int_0^t e_{r(0)}e^{W_{00}s}W_{01}e^{W_{11}(t-s)}V^{-1}\mathbb{1}\mathrm{d}s\right] \\
&= \sum_k \mathbb{E}c(0)^2I_{\{r(0)=k\}}v_ke_ke^{W_{11}t}V^{-1}\mathbb{1}+\int_0^t \mathbb{E}c(0)I_{\{r(0)=k\}}e_ke^{W_{00}s}W_{01}e^{W_{11}(t-s)}V^{-1}\mathbb{1}\mathrm{d}s \\
&= \sum_k \mu_{2k}v_ke_ke^{W_{11}t}V^{-1}\mathbb{1}+\int_0^t \mu_{1k}e_ke^{W_{00}s}W_{01}e^{W_{11}(t-s)}V^{-1}\mathbb{1}\mathrm{d}s \\
&= \mu_2Ve^{W_{11}t}V^{-1}\mathbb{1}+\int_0^t \mu_1e^{W_{00}s}W_{01}e^{W_{11}(t-s)}V^{-1}\mathbb{1}\mathrm{d}s \\
&= (m_1+m_2)V^{-1}e^{W_{11}t}V^{-1}\mathbb{1}+\int_0^t m_1V^{-1}e^{W_{00}s}W_{01}e^{W_{11}(t-s)}V^{-1}\mathbb{1}\mathrm{d}s,
\end{aligned}
$$

where $m_1$ and $m_2$ are the steady-state values of moments given in Eq. (4). Since the autocorrelation function is defined as $R_c(t)=\mathbb{E}c(0)c(t)-\mathbb{E}c(0)\mathbb{E}c(t)$, we finally obtain the explicit expression of the autocorrelation function, which is given by

$$
R_c(t) = (m_1+m_2)V^{-1}e^{W_{11}t}V^{-1}\mathbb{1}+\int_0^t m_1V^{-1}e^{W_{00}s}W_{01}e^{W_{11}(t-s)}V^{-1}\mathbb{1}\mathrm{d}s-(m_1V^{-1}\mathbb{1})^2. \quad (23)
$$

Here the autocorrelation function is expressed in matrix form. A more explicit expression can be obtained by expanding the matrix exponentials $e^{W_{11}t}$ and $e^{W_{00}s}$ in terms of their eigenvalues and eigenvectors. For simplicity, we next focus on the timer strategy. The results for other control strategies can be obtained from the results for the timer strategies by substituting the parameters $N$ and $N_0$ for the effective parameters $\bar{N}$ and $\bar{N}_0$, respectively (see Sec. 1.4 for details). With these notation in Sec. 1.4, the autocorrelation function given in Eq. (23) can be rewritten as

$$
R_c(t) = \frac{1}{v_1^2}\left[(m_1+m_2)M^2Re^{D_{11}t}\bar{R}'\mathbb{1}+\int_0^t m_1MRe^{D_{00}s}\bar{R}'W_{01}MRe^{D_{11}(t-s)}\bar{R}'\mathbb{1}\mathrm{d}s-(m_1M\mathbb{1})^2\right].
$$

This suggests that the autocorrelation function (power spectrum) is the linear combination of $2N-1$ exponential (Lorentzian) functions:

$$
R(t)=\sum_{k=2}^{2N}u_ke^{\lambda_kt},\quad G(\xi)=\sum_{k=2}^{2N}\frac{-2u_k\lambda_k}{4\pi^2\xi^2+\lambda_k^2},
$$

where $\lambda_1,\cdots,\lambda_{2N}$ are all the eigenvalues of $W_{00}$ and $W_{11}$, all the coefficients $u_k$ associated with the eigenvalues of $W_{00}$ are given by

$$
u_k=\frac{1}{v_1^2}\sum_{j=1}^N\frac{[m_1MR]_k[\bar{R}'W_{01}MR]_{kj}[\bar{R}'\mathbb{1}]_j}{\lambda_k-\lambda_{N+j}},\quad 2\le k\le N,
$$

and all the coefficients $u_k$ associated with the eigenvalues of $W_{11}$ are given by

$$
u_{N+k}=\frac{1}{v_1^2}[(m_1+m_2)M^2R]_k[\bar{R}'\mathbb{1}]_k-\frac{1}{v_1^2}\sum_{j=1}^N\frac{[m_1MR]_j[\bar{R}'W_{01}MR]_{jk}[\bar{R}'\mathbb{1}]_k}{\lambda_j-\lambda_{N+k}},\quad 1\le k\le N.
$$

Here $[m]_k$ denotes the $k$th entry of the vector $m$ and $A_{kl}$ denotes the $(k,j)$-th entry of the matrix $A$. In fact, the coefficients $u_k$ can be computed more explicitly. To see this, note that

$$[\bar{R}'\mathbb{1}]_k = \frac{1}{\sqrt{N}}\sum_{j=1}^{N}[\bar{R}]_{jk} = \frac{1}{\sqrt{N}}\sum_{j=1}^{N}\bar{\omega}_k^{j-1} = \sqrt{N}\delta_{k1}.$$

Thus all the coefficients $u_k$ associated with the eigenvalues of $W_{00}$ can be simplified as

$$u_k = \frac{\sqrt{N}}{v_1^2}\frac{[m_1MR]_k[\bar{R}'W_{01}MR]_{k1}}{\lambda_k - \lambda_{N+1}}, \quad 1 \leq k \leq N, \tag{24}$$

and all the coefficients $u_k$ associated with the eigenvalues of $W_{11}$ can be simplified as

$$\begin{aligned} &u_{N+k} = 0, \quad 2 \leq k \leq N, \\ &u_{N+1} = \frac{\sqrt{N}}{v_1^2}[(m_1+m_2)M^2R]_1 - \frac{\sqrt{N}}{v_1^2}\sum_{j=1}^{N}\frac{[m_1MR]_j[\bar{R}'W_{01}MR]_{j1}}{\lambda_j - \lambda_{N+1}}. \end{aligned} \tag{25}$$

This suggests that the autocorrelation function (power spectrum) is actually the linear combination of only $N$ exponential (Lorentzian) functions:

$$R_c(t) = \sum_{k=2}^{N+1}u_ke^{\lambda_kt}, \quad G_c(\xi) = \sum_{k=2}^{N+1}\frac{-2u_k\lambda_k}{4\pi^2\xi^2+\lambda_k^2},$$

Combining Eqs. (4), (13), and (15), we have

$$m_1MR = -m_0W_{01}W_{11}^{-1}MR = -\frac{1}{N}\mathbb{1}'W_{01}MRD_{11}^{-1}.$$

It is easy to check that

$$W_{01} = SH = \rho Bv_1^{\beta}\text{diag}(1,\cdots,2^{\beta(N_0-1)/N},\kappa2^{\beta N_0/N},\cdots,\kappa2^{\beta(N-1)/N}).$$

This shows that

$$\begin{aligned} [m_1MR]_k &= -\frac{1}{N}\sum_{j=1}^{N}[W_{01}M]_{jj}R_{jk}[D_{11}^{-1}]_{kk} = -\frac{1}{N^{3/2}\lambda_{N+k}}\sum_{j=1}^{N}[W_{01}M]_{jj}\omega_k^{j-1} \\ &= \frac{\rho Bv_1^{\beta}}{N^{3/2}(d+a-2^{-1/N}a\omega_k)}\left[\sum_{j=1}^{N_0}2^{(\beta-1)(j-1)/N}\omega_k^{j-1} + \kappa\sum_{j=N_0+1}^{N}2^{(\beta-1)(j-1)/N}\omega_k^{j-1}\right] \\ &= \frac{\rho Bv_1^{\beta}}{N^{3/2}(d+a-2^{-1/N}a\omega_k)}\left[\sum_{j=1}^{N_0}(2^{(\beta-1)/N}\omega_k)^{j-1} + \kappa\sum_{j=N_0+1}^{N}(2^{(\beta-1)/N}\omega_k)^{j-1}\right] \\ &= \frac{\rho Bv_1^{\beta}\Delta_k}{N^{3/2}(d+a-2^{-1/N}a\omega_k)}, \end{aligned}$$

where

$$\begin{aligned} \Delta_k &= \sum_{j=1}^{N_0}(2^{(\beta-1)/N}\omega_k)^{j-1} + \kappa\sum_{j=N_0+1}^{N}(2^{(\beta-1)/N}\omega_k)^{j-1} \\ &= \frac{1-\kappa2^{(\beta-1)}+(\kappa-1)2^{(\beta-1)w}\omega_k^{N_0}}{1-2^{(\beta-1)/N}\omega_k}, \end{aligned}$$

with $w = N_0/N$ being the proportion of cell cycle before replication. Moreover, we have

$$[\bar{R}'W_{01}MR]_{k1} = \sum_{j=1}^{N}[\bar{R}]_{jk}[W_{01}M]_{jj}[R]_{j1} = \frac{1}{N}\sum_{j=1}^{N}\bar{\omega}_k^{j-1}[W_{01}M]_{jj} = \frac{\rho B v_1^{\beta}\bar{\Delta}_k}{N}.$$

It then follows from Eq. (24) that

$$u_k = \frac{\rho^2 B^2 v_1^{2\beta-2}|\Delta_k|^2}{N^2(d+a-2^{-1/N}a\omega_k)(d+a\omega_k-2^{-1/N}a)}, \quad 1 \leq k \leq N.$$

In addition, it is easy to see that

$$[(m_1+m_2)M^2R]_1 = \sum_{j=1}^{N}[(m_1+m_2)M^2]_j R_{j1} = \frac{1}{\sqrt{N}}(m_1+m_2)M^2\mathbb{1}.$$

It thus follows from Eq. (25) that

$$u_{N+1} = \frac{1}{v_1^2}(m_1+m_2)M^2\mathbb{1} - \sum_{k=1}^{N}u_k = \langle c(t)\rangle^2 - \sum_{k=1}^{N}u_k.$$

To proceed, note that the concentration mean can be computed explicitly as

$$\begin{aligned}
\langle c(t)\rangle &= m_1V^{-1}\mathbb{1} = -\frac{1}{Nv_1}\mathbb{1}'W_{01}W_{11}^{-1}M\mathbb{1} = -\frac{1}{Nv_1}\mathbb{1}'W_{01}MRD_{11}^{-1}\bar{R}'\mathbb{1} \\
&= \frac{1}{Nv_1}\sum_{j,k=1}^{N}\frac{[W_{01}M]_{jj}R_{jk}[\bar{R}'\mathbb{1}]_k}{\lambda_{N+k}} \\
&= \frac{1}{Nv_1(d+a-2^{-1/N}a)}\sum_{j=1}^{N}[W_{01}M]_{jj} \\
&= \frac{\rho B v_1^{\beta-1}\Delta_1}{N(d+a-2^{-1/N}a)} = \sqrt{u_1}.
\end{aligned}$$

Thus we obtain

$$u_{N+1} = \langle c(t)\rangle^2 - \langle c(t)\rangle^2 - \sum_{k=2}^{N}u_k = \sigma_c^2 - \sum_{k=2}^{N}u_k,$$

where $\sigma_c^2$ is the steady-state variance of gene product concentration.

In summary, we have proved that the autocorrelation function (power spectrum) is the weighted sum of $N$ exponential (Lorentzian) functions:

$$R_c(t) = \sum_{k=1}^{N}u_ke^{\lambda_k t}, \quad G_c(\xi) = \sum_{k=1}^{N}\frac{-2u_k\lambda_k}{4\pi^2\xi^2+\lambda_k^2}, \tag{26}$$

where the exponents $\lambda_k$ are given by

$$\begin{aligned}
&\lambda_k = -a(1-\omega_k), \quad 1 \leq k \leq N-1, \\
&\lambda_N = -d-a+2^{-1/N}a = -d-a(1-e^{-\log(2)/N}) \approx -d-\log(2)f = -d_{\text{eff}},
\end{aligned}$$

with $\omega_k = e^{2k\pi i/N}$ being all $N$th roots of unity, and the coefficients $u_k$ are given by

$$\begin{aligned}
&u_k = \frac{\rho^2 B^2 v_1^{2\beta-2}|\Delta_k|^2}{N^2(d+a-2^{-1/N}a\omega_k)(d+a\omega_k-2^{-1/N}a)}, \quad 1 \leq k \leq N-1, \\
&u_N = \sigma_c^2 - \sum_{k=1}^{N-1}u_k,
\end{aligned}$$

with $\sigma_c^2$ being the steady-state variance of concentration and with $\Delta_k$ being defined as

$$\Delta_k = \frac{1 - \kappa 2^{(\beta-1)} + (\kappa - 1)2^{(\beta-1)w}\omega_k^{N_0}}{1 - 2^{(\beta-1)/N}\omega_k}. \tag{27}$$

In the special case of perfect concentration homeostasis ($\beta = \kappa = 1$), we have $\Delta_k = u_k = 0$ for each $1 \leq k \leq N-1$ and thus the autocorrelation function (power spectrum) reduces to the following exponential (Lorentzian) function:

$$R_c(t) = \sigma_c^2 e^{-d_{\text{eff}}t}, \quad G_c(\xi) = \frac{-2\sigma_c^2 d_{\text{eff}}}{4\pi^2\xi^2 + d_{\text{eff}}^2}.$$

In this case, both the autocorrelation function and power spectrum are monotonic decreasing functions and thus no concentration oscillations can be observed.

If homeostasis is not perfect, the power spectrum for concentration can either be monotonically decreasing or have an off-zero peak. When $N \gg 1$, the position of the off-zero peak is equal to the cell cycle frequency $f$, the width of the off-zero peak is given by $D = 2\pi f/N$. The absolute height of the zero peak is given by

$$H_{\text{zero}} = G_c(0) = -2\sum_{k=1}^{N}\frac{u_k}{\lambda_k}.$$

Moreover, the absolute height of the off-zero peak is given by

$$H_{\text{off}-\text{zero}} = \frac{-2u_1\lambda_1}{4\pi^2 f^2 + \lambda_1^2} + \frac{-2u_1\lambda_1}{4\pi^2 f^2 + \lambda_1^2} = -4\text{Re}\left(\frac{u_1\lambda_1}{4\pi^2 f^2 + \lambda_1^2}\right),$$

where $\text{Re}(z)$ denotes the real part of $z$. Since we have normalized the power spectrum so that $G_c(0) = 1$, the height of the off-zero peak is then the ratio of the absolute heights of the off-zero and zero peaks, which is given by

$$H = \frac{H_{\text{zero}}}{H_{\text{off}-\text{zero}}} = \frac{2\text{Re}(u_1\lambda_1/(4\pi^2 f^2 + \lambda_1^2))}{\sum_{k=1}^{N} u_k/\lambda_k}. \tag{28}$$

Note that the height $H$ is proportional to $u_1$, which is proportional to $|\Delta_1|^2$. It follows from Eq. (27) that $|\Delta_1|^2$ is proportional to

$$C_1(w,\kappa,\beta) = |1 - \kappa 2^{(\beta-1)} + (\kappa-1)2^{(\beta-1)w}\omega_1^{N_0}|^2 = |1 - \kappa 2^{(\beta-1)} + (\kappa-1)2^{(\beta-1)w}e^{2\pi wi}|^2.$$

Therefore, the height $H$ is also proportional to $C_1(w,\kappa,\beta)$ and thus vanishes if $C_1(w,\kappa,\beta) = 0$.

##### 3.3 Height of the off-zero peak for unstable gene products

Here we focus on the power spectrum for unstable gene products. Without loss of generality, we assume that at least one of $\beta$ and $\kappa$ is not equal to 1. For unstable gene products, we have $d \gg a$ and thus $|\lambda_{N+1}| \gg |\lambda_k|$ for $2 \leq k \leq N$. Thus the power spectrum of gene product concentration is approximately given by

$$G_c(\xi) = \sum_{k=1}^{N-1}\frac{-2u_k\lambda_k}{4\pi^2\xi^2 + \lambda_k^2},$$

where $\lambda_k = -a + a\omega_k$ with $\omega_k = e^{2k\pi i/N}$ and

$$u_k = \frac{\rho^2 B^2 v_1^{2\beta-2}|\Delta_k|^2}{a^2\eta^2}.$$

Note that the power spectrum can be rewritten as

$$G_c(\xi) = \frac{2\rho^2B^2v_1^{2\beta-2}}{a\eta^2}\sum_{k=1}^{N-1}\frac{|\Delta_k|^2(1-\omega_k)}{4\pi^2\xi^2+a^2(1-\omega_k)^2}.$$

When $N \gg 1$, the power spectrum has the following approximation:

$$G_c(\xi) \approx \frac{2\rho^2B^2v_1^{2\beta-2}}{a\eta^2}\sum_{k=1}^{[N/2]}|\Delta_k|^2G_k(\xi),$$

where

$$G_k(\xi) = \frac{1-\omega_k}{4\pi^2\xi^2+a^2(1-\omega_k)^2} + \frac{1-\bar{\omega}_k}{4\pi^2\xi^2+a^2(1-\bar{\omega}_k)^2}.$$

Straightforward computations show that

$$G_k(\xi) = \frac{\sin^2\theta_k(\pi^2\xi^2+a^2\sin^2\theta_k)}{(\pi^2\xi^2-a^2\sin^2\theta_k)^2+4\pi^2\xi^2a^2\sin^4\theta_k},$$

where $\theta_k = k\pi/N$. In fact, the function $G_k(\xi)$ characterizes the $k$th off-zero peak. It is easy to check that the position of the $k$th off-zero peak is given by

$$\xi = \frac{a}{\pi}\sin\theta_k\sqrt{2\cos\theta_k - 1}.$$

When $N \gg 1$, we have $\sin\theta_k \approx \theta_k$ and $\cos\theta_k \approx 1$ and thus the position of the $k$th off-zero peak is approximately given by

$$\xi = \frac{a\theta_k}{\pi} \approx kf,$$

and the function $G_k(\xi)$ can be further simplified as

$$G_k(\xi) \approx \frac{k^2(\xi^2+k^2f^2)}{N^2(\xi^2-k^2f^2)^2+4k^4\pi^2f^2\xi^2}. \tag{29}$$

In addition, we have

$$|\Delta_k|^2 = \frac{|1-\kappa 2^{(\beta-1)}+(\kappa-1)2^{(\beta-1)w}\omega_k^{N_0}|^2}{|1-2^{(\beta-1)/N}\omega_k|^2} = \frac{C_k(w,\kappa,\beta)}{|1-2^{(\beta-1)/N}\omega_k|^2},$$

where

$$C_k(w,\kappa,\beta) = 2^{2(\beta-1)w}(\kappa-1)^2+(1-\kappa 2^{\beta-1})^2+2^{(\beta-1)w+1}(\kappa-1)(1-\kappa 2^{\beta-1})\cos(2k\pi w)$$

is a function of $w$, $\kappa$, and $\beta$. Moreover, note that

$$\begin{aligned}|1-2^{(\beta-1)/N}\omega_k|^2 &= (1-2^{(\beta-1)/N})^2+2^{(\beta-1)/N+2}\sin^2\theta_k \\ &\approx \frac{4k^2\pi^2+(\log 2)^2(\beta-1)^2}{N^2}.\end{aligned}$$

Thus we have

$$|\Delta_k|^2 \approx \frac{C_k(w,\kappa,\beta)N^2}{4k^2\pi^2+(\log 2)^2(\beta-1)^2}.$$

Thus when $N \gg 1$, the power spectrum for unstable gene products can be simplified as

$$G_c(\xi) \approx \frac{2\rho^2B^2v_1^{2\beta-2}N}{f\eta^2}\sum_{k=1}^{\infty}\frac{C_k(w,\kappa,\beta)}{4k^2\pi^2+(\log 2)^2(\beta-1)^2}G_k(\xi),$$

where $G_k(\xi)$ is given in Eq. (29). Note that the maximum of $G_k(\xi)$ is approximately given by

$$G_k(kf) \approx \frac{2k^4 f^2}{4k^6\pi^2 f^4} = \frac{1}{2k^2\pi^2 f^2}.$$

Thus the absolute height of the off-zero peak is given by

$$H_{\text{off-zero}} \approx \frac{2\rho^2 B^2 v_1^{2\beta-2} N}{f\eta^2} \times \frac{C_1(w,\kappa,\beta)}{4\pi^2 + (\log 2)^2(\beta-1)^2} \times \frac{1}{2\pi^2 f^2}.$$

Moreover, the absolute height of the zero peak is given by

$$\begin{aligned} H_{\text{zero}} &\approx \frac{2\rho^2 B^2 v_1^{2\beta-2} N}{f\eta^2} \sum_{k=1}^{\infty} \frac{C_k(w,\kappa,\beta)}{4k^2\pi^2 + (\log 2)^2(\beta-1)^2} G_r(0) \\ &= \frac{2\rho^2 B^2 v_1^{2\beta-2} N}{f\eta^2} \times \frac{1}{N^2 f^2} \times \sum_{k=1}^{\infty} \frac{C_k(w,\kappa,\beta)}{4k^2\pi^2 + (\log 2)^2(\beta-1)^2}. \end{aligned}$$

Since we have normalized the power spectrum so that $G_c(0) = 1$, the height of the off-zero peak is then the ratio of the absolute heights of the off-zero and zero peaks, which is given by

$$H = \frac{H_{\text{off-zero}}}{H_{\text{zero}}} \approx \frac{C_1(w,\kappa,\beta) N^2}{2\pi^2(4\pi^2 + (\log 2)^2(\beta-1)^2) \sum_{k=1}^{\infty} \frac{C_k(w,\kappa,\beta)}{4k^2\pi^2+(\log 2)^2(\beta-1)^2}}.$$

Since $(\log 2)^2(\beta-1)^2 \ll 4\pi^2$, the height of the off-zero peak can be simplified as

$$H \approx \frac{C_1(w,\kappa,\beta) N^2}{2\pi^2 C(w,\kappa,\beta)},$$

where

$$C(w,\kappa,\beta) = \sum_{k=1}^{\infty} \frac{C_k(w,\kappa,\beta)}{k^2}.$$

Therefore, the height $H$ is also proportional to $N^2$. We emphasize that while this conclusion is derived for unstable gene products, it also holds for all gene products, according to our simulations.

#### Note 4. Technical details for Fig. 3

In (a)-(c), the model parameters are chosen as $N = 50, B = 1, \alpha_0 = \alpha_1 = 1, d = \eta f$. The parameters $\rho$ and $a$ are chosen so that $\langle n \rangle = 100$ and $\langle V \rangle = 1$. The remaining parameters are chosen as $N_0 = 25, \eta = 1$ for (a), $N_0 = 25, \kappa = 2$ for (b) and $\beta = 0, \eta = 1$ for (c). While we have assumed the adder strategy in the simulations, similar results also hold for other control strategies.

In (d)-(f), the model parameters are chosen as $N_0 = 0.5N, B = 1, \alpha_0 = \alpha_1 = 0.01, d = \eta f$. The parameters $\rho$ and $a$ are chosen so that $\langle n \rangle = 500$ and $\langle V \rangle = 100$. The remaining parameters are chosen as $N = 50, \beta = \kappa = 1, \eta = 0.2$ for the left panel, $N = 50, \beta = 0, \kappa = \sqrt{2}, \eta = 0.5$ for the middle panel, and $N = 140, \beta = 1, \kappa = 2, \eta = 2$ for the right panel.

In (a)-(f), the growth rate $g$ is determined so that $f = 0.1$.

#### Note 5. Technical details for Fig. 4

In (e), the model parameters are chosen as $N = 50, N_0 = 23, \rho = 17, B = 1, \beta = 1, \kappa = 2, d = 0.1, \eta = 1$. The parameter $a$ is chosen so that $\langle V \rangle = 1$. The strengths $\alpha_0$ and $\alpha_1$ of size control are chosen as $\alpha_0 = \alpha_1 = 1$ for the upper panel (adder), $\alpha_0 = 0.5, \alpha_1 = 2$ for the middle panel (timer-sizer), and $\alpha_0 = 2, \alpha_1 = 0.5$ for the lower panel (sizer-timer). The growth rate $g$ is determined so that $f = 0.1$.

#### Note 6. Technical details for Fig. 5

In (a)-(c), the model parameters are chosen as $N = 50, B = 1, \kappa = 2, \alpha_0 = \alpha_1 = 1, d = \eta f$. The parameters $\rho$ and $a$ are chosen so that $\langle n \rangle = 100$ and $\langle V \rangle = 1$. The remaining parameters are chosen as $N_0 = 25, \beta = 0, \eta = 0$ for (a), $N_0 = 10, \beta = 0, \eta = 3$ for (b), and $N_0 = 16, \beta = 1, \eta = 10$ for (c).

In (d), the model parameters are chosen as $N = 30, N_0 = 11, \rho = 66, B = 1, \beta = 1, \kappa = 2, d = 1, \eta = 10$. The parameter $a$ is chosen so that $\langle V \rangle = 1$. The strengths $\alpha_0$ and $\alpha_1$ of size control are chosen as $\alpha_0 = \alpha_1 = 1$ for the blue curve, $\alpha_0 = 0.5, \alpha_1 = 2$ for the red curve, and $\alpha_0 = 2, \alpha_1 = 0.5$ for the green curve.

In (e),(f), the model parameters are chosen as $N = 30, N_0 = 18, \alpha_0 = \alpha_1 = 1, d = \eta f$. The parameters $\rho$ and $a$ are chosen so that $\langle n \rangle = 100$ and $\langle V \rangle = 1$. The remaining parameters are chosen as $B = 0.2, \eta = 10$ for (e) and $\beta = 1, \kappa = 2$ for (f).

In (a)-(f), the growth rate $g$ is determined so that $f = 0.1$.

#### References

[1] Jia, C., Singh, A. & Grima, R. Cell size distribution of lineage data: analytic results and parameter inference. *iScience* **24**, 102220 (2021).

[2] Paulsson, J. & Ehrenberg, M. Random signal fluctuations can reduce random fluctuations in regulated components of chemical regulatory networks. *Phys. Rev. Lett.* **84**, 5447 (2000).

[3] Jia, C., Zhang, M. Q. & Hong, Q. Emergent Lévy behavior in single-cell stochastic gene expression. *Phys. Rev. E* **96**, 040402(R) (2017).

[4] Jia, C., Yin, G. G., Zhang, M. Q. *et al.* Single-cell stochastic gene expression kinetics with coupled positive-plus-negative feedback. *Phys. Rev. E* **100**, 052406 (2019).

[5] Friedman, N., Cai, L. & Xie, X. S. Linking stochastic dynamics to population distribution: an analytical framework of gene expression. *Phys. Rev. Lett.* **97**, 168302 (2006).

[6] Claude, K.-L. *et al.* Transcription coordinates histone amounts and genome content. *Nat. Commun.* **12**, 1–17 (2021).

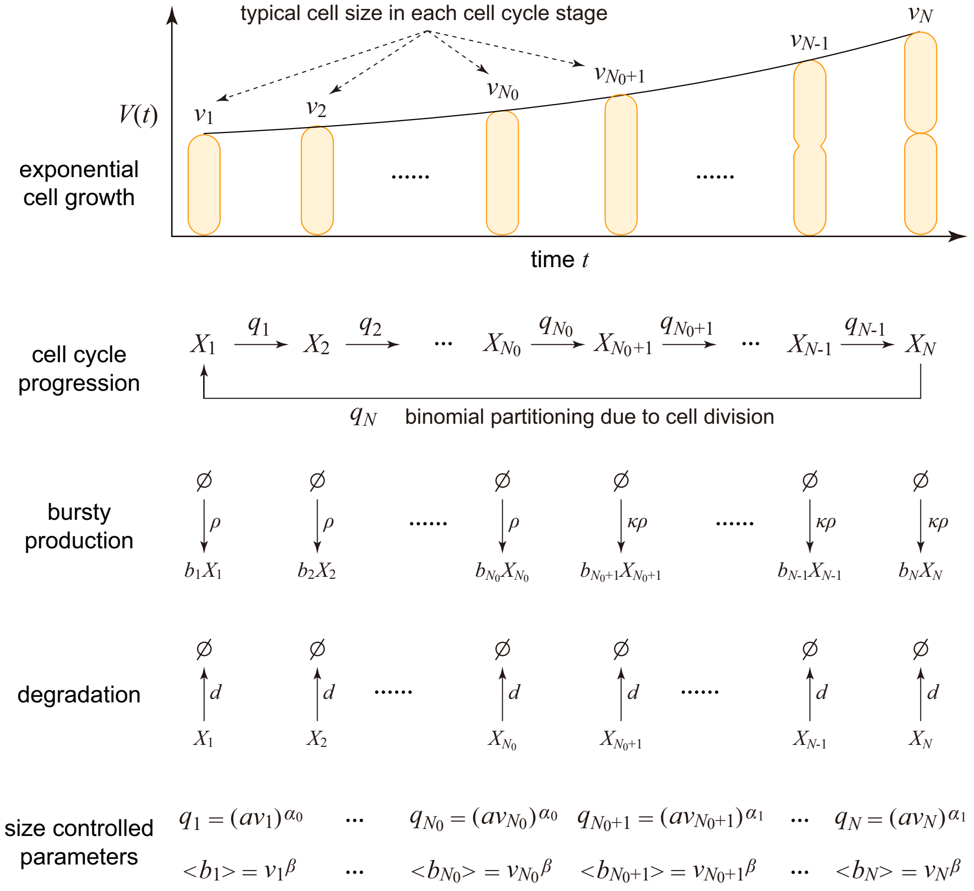

Fig. 1: **Reduced model obtained by making the mean-field approximation of cell size dynamics.** Here $X_k$ denotes the gene product at stage $k$ and $b_k$ denotes the burst size at stage $k$. Moreover, $v_k$ is the typical cell size at stage $k$, $B_k$ is the mean burst size at stage $k$, and $q_k$ is the typical transition rate from stage $k$ to the next.

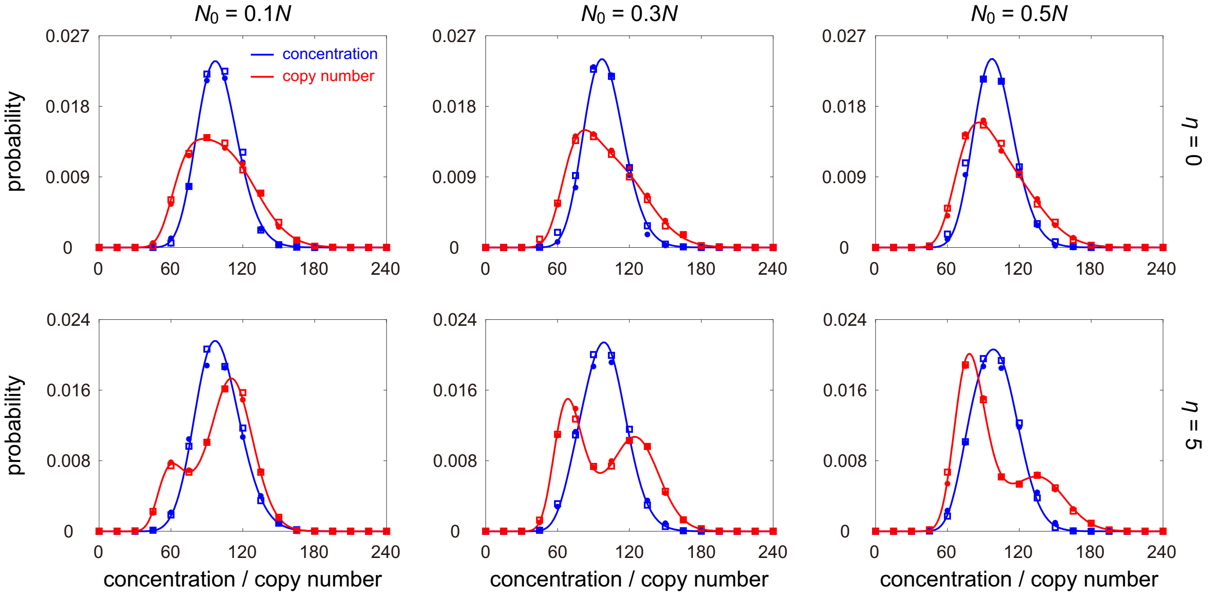

Fig. 2: **Comparison between the full and mean-field models as $N_0$ and $\eta$ vary.** The blue (red) dots show the simulated concentration (copy number) distribution for the full model obtained from the stochastic simulation algorithm (SSA). The blue (red) squares show the simulated concentration (copy number) distribution for the mean-field model obtained from SSA. The blue (red) curve shows the analytical approximate concentration (copy number) distribution given in Eq. (4) (Eq. (13)) in the main text. The model parameters are chosen as $N = 50, B = 1, \beta = 0, \kappa = 2, \alpha_0 = \alpha_1 = 1, d = \eta f$. The growth rate $g$ is determined so that $f = 0.1$. The parameter $\rho$ and $a$ are chosen so that $\langle n \rangle = 100$ and $\langle V \rangle = 1$.

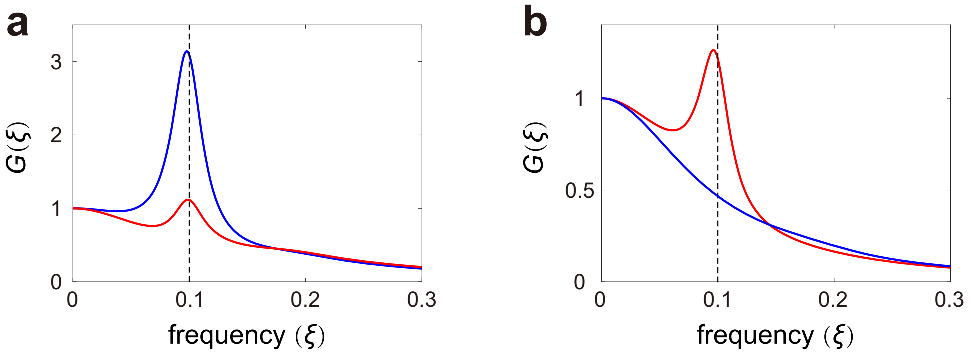

Fig. 3: **Power spectra of copy number and concentration fluctuations.** **(a)** Power spectra of copy number fluctuations when $w = 0.5$ (blue curve) and $w = 0.8$ (red curve). **(b)** Power spectra of concentration fluctuations when $w = 0.5$ (blue curve) and $w = 0.8$ (red curve). In (a),(b), the model parameters are chosen as $N = 20, B = 1, \beta = 0, \kappa = \sqrt{2}, \alpha_0 = \alpha_1 = 0.01, d = 0.5, \eta = 5$. The growth rate $g$ is determined so that $f = 0.1$. The parameter $\rho$ and $a$ are chosen so that $\langle n \rangle = 100$ and $\langle V \rangle = 1$. The power spectra are computed using Eq. (26).

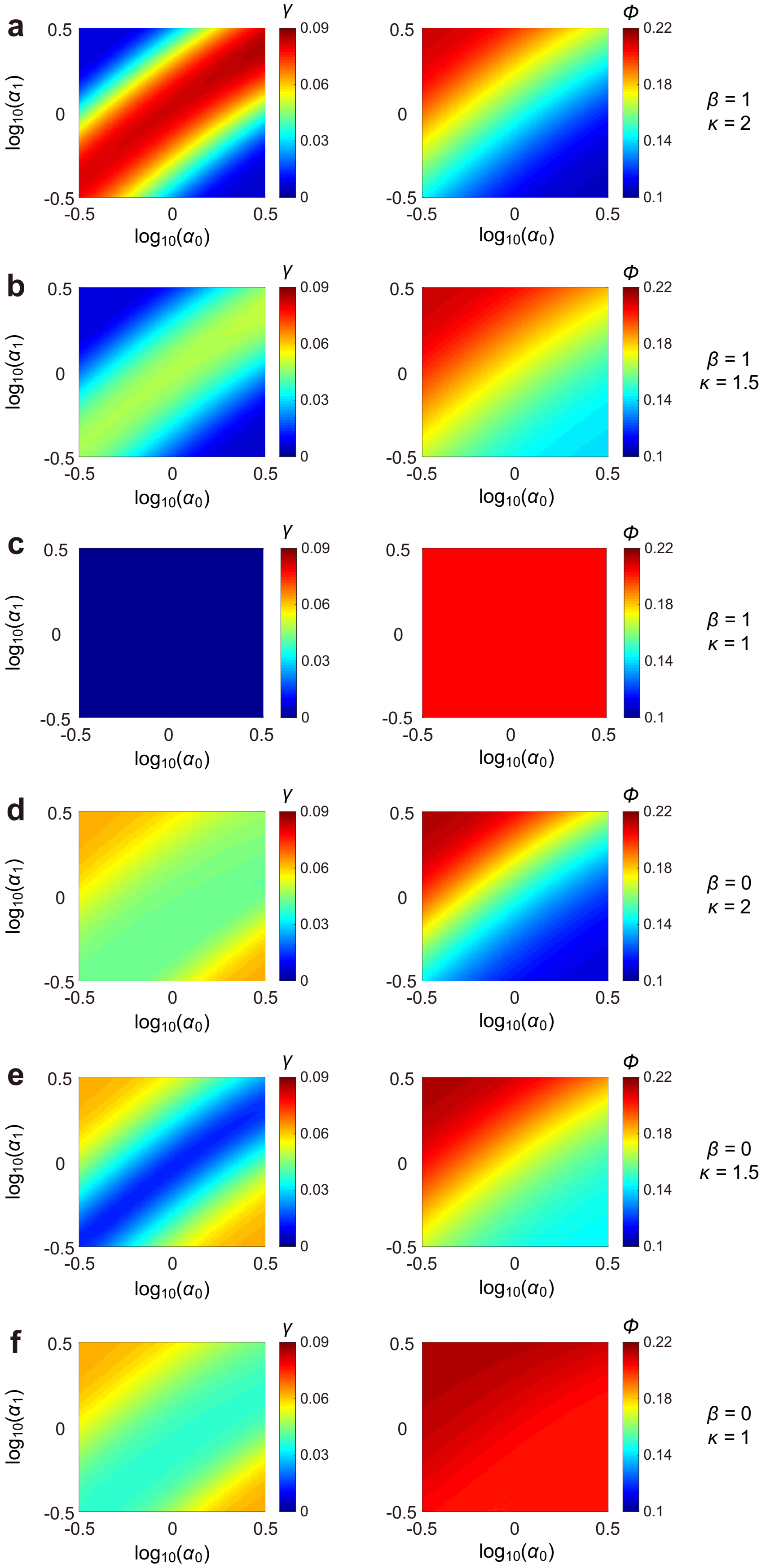

Fig. 4: **Influence of size control on concentration homeostasis and concentration noise for various choices of $\beta$ and $\kappa$.** (**a**)-(**f**) Heat plot of the homeostasis accuracy $\gamma$ (left panel) and concentration noise $\phi$ (right panel) versus the strengths of size control, $\alpha_0$ and $\alpha_1$. The model parameters are chosen as $N = 50, N_0 = 23, \rho = 1.7, B = 1, d = 0.1, \eta = 1$. In (c), perfect concentration homeostasis is present and thus both $\gamma$ and $\phi$ are almost independent of $\alpha_0$ and $\alpha_1$.

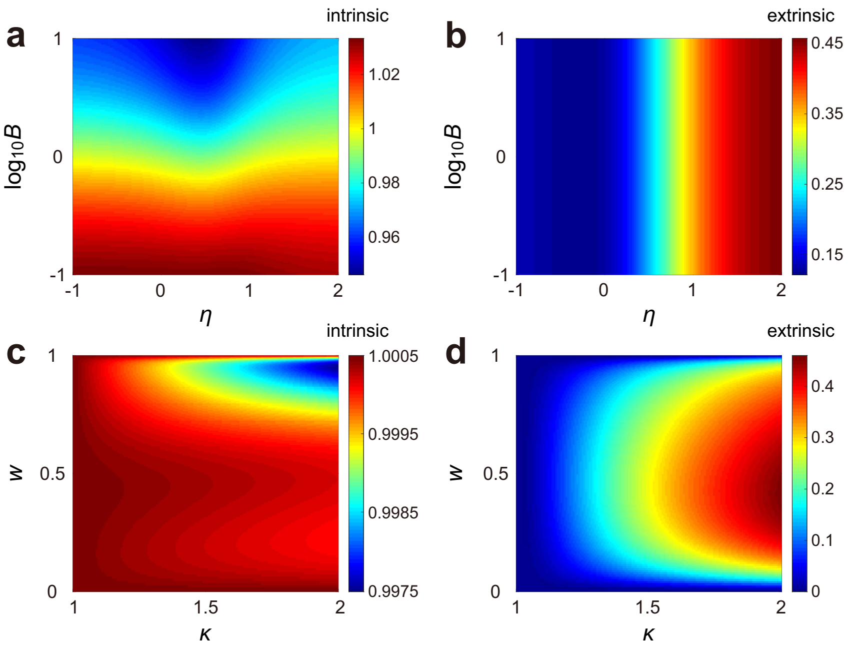

Fig. 5: **Comparison between the concentration and copy number noise when the burst size scales with cell size. (a)** Ratio of the intrinsic parts of concentration noise and copy number noise versus $\eta$ and $B$. **(b)** Ratio of the extrinsic parts of concentration noise and copy number noise versus $\eta$ and $B$. **(c)** Ratio of the intrinsic parts of concentration noise and copy number noise versus $\kappa$ and $w$. **(d)** Ratio of the extrinsic parts of concentration noise and copy number noise versus $\kappa$ and $w$. In the presence of balanced biosynthesis ($\beta = 1$), intrinsic noise in concentration is almost the same as that in copy number, while extrinsic noise in concentration is always smaller than that in copy number. Extrinsic noise in concentration is smaller than that in the copy number by at least 53%, where this lower bound is obtained when $\kappa = 2$, $w \approx 0.4$, and $\eta \gg 1$. In (a),(b), the model parameters are chosen as $N = 50, N_0 = 25, \beta = 1, \kappa = 2, \alpha_0 = \alpha_1 = 1, d = \eta f$. In (c),(d), the model parameters are chosen as $N = 50, B = 1, \beta = 1, \alpha_0 = \alpha_1 = 1, d = 10, \eta = 100$. In (a)-(d), the growth rate $g$ is determined so that $f = 0.1$ and the parameters $\rho$ and $a$ are chosen so that $\langle n \rangle = 100$ and $\langle V \rangle = 1$. The intrinsic and extrinsic parts of gene expression noise are computed using Eqs. (8) and (9).

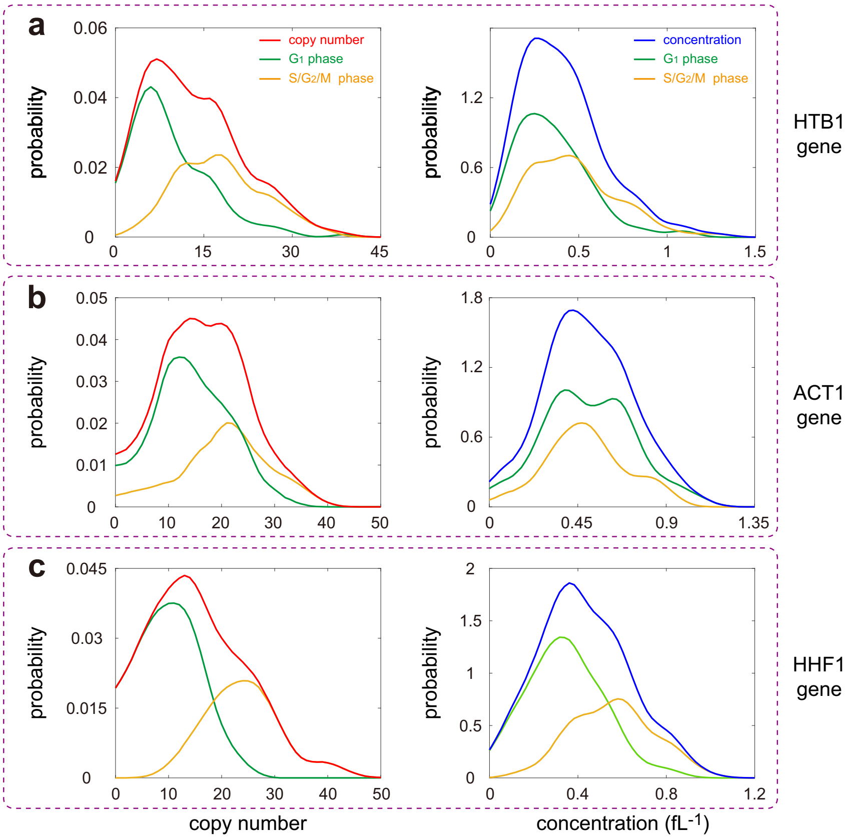

Fig. 6: **Copy number and concentration distributions of mRNAs in budding yeast cells. (a)** The mRNA copy number (red curve) and concentration (blue curve) distributions for the *HTB1* gene. The green curves show the distributions of cells in the $G_1$ phase and the yellow curves show the distributions of cells in the $G_2$/S/M phase. **(b)** Same as (a) but for the *ACT1* gene. **(c)** Same as (a) but for the *HHF1* gene. The data shown are published in [6].
